# The SUMO activating enzyme subunit, SAE2, contributes SUMO protein bias for mitotic fidelity

**DOI:** 10.1101/2022.03.18.484840

**Authors:** Alexandra K. Walker, Alexander J. Lanz, Mohammed Jamshad, Alexander J. Garvin, Peter Wotherspoon, Benjamin F. Cooper, Timothy J. Knowles, Joanna R. Morris

## Abstract

Mammalian cells possess three conjugatable SUMO variants: SUMO1 and the largely indistinguishable SUMO2 and SUMO3 (designated SUMO2/3). Some SUMOylated substrates are modified by both SUMO1 and SUMO2/3, while others show biased modifications towards SUMO1 or SUMO2/3. How preferential SUMO protein conjugation is coordinated is poorly understood.

Here, we examine a modification of the catalytic component of the human SUMO Activation Enzyme, SAE2. We observe that lysine 164 of SAE2 is deacetylated during mitosis in an HDAC6-dependent manner. We find that an acetyl-analogue mutant, SAE2-K164Q, biases the activation and conjugation of SUMO2>SUMO1 and discriminates SUMO1 and SUMO2/3 through their C-terminal tails.

Complementation of SAE2-depleted or inhibited cells with SAE2-K164Q restricts mitotic SUMO1-conjugates and increases multipolar spindle formation. We confirm the SUMO E1-dependent modification of the nuclear mitotic apparatus, NuMA, and find that the mitotic defects of both SAE2-K164Q complemented cells and HDAC6-inhibitor-treated cells are corrected by either over-expression of SUMO1 or by expression of a GFP-SUMO1-NuMA-K1766R fusion protein.

Our observations suggest a model in which the SAE1:SAE2 enzyme is deacetylated on early mitosis to encourage the conjugation of SUMO1 to support mitotic fidelity. These surprising data reveal that the SUMO-activating enzyme can bias SUMO variant conjugation.

## Introduction

Mammals have ∼20 ubiquitin-like modifiers, including the Small Ubiquitin-like Modifiers (SUMOs) [1]. Higher eukaryotes express two subfamilies of conjugatable SUMO variants: SUMO1 and the highly similar SUMO2 and SUMO3, referred to as SUMO2/3. SUMOylation contributes to many intracellular processes, including transcription, DNA repair, chromatin remodelling, signal transduction, and mitosis [2, 3]. In the initial step of the SUMOylation cascade, SUMO activation is catalysed by the sole SUMO E1 heterodimer comprising SUMO-Activating Enzyme 1 (SAE1/AOS1) and 2 (SAE2/UBA2) [4–6]; herein referred to as SAE1:SAE2. The SAE1:SAE2 adenylation catalytic site coordinates SUMO and ATP-Mg^2+^ and initiates the conjugation process by adenylating the SUMO C-terminus, producing SUMO-AMP. Subsequently, SAE1:SAE2 undergoes remodelling around SUMO-AMP to form the thiolation catalytic site for thioester bond formation between the SUMO C-terminus and SAE2-C173 [7]. The activated SUMO is then transferred to the catalytic cysteine in the only SUMO-conjugating enzyme, UBC9 [8]. SUMO can then be transferred to a target lysine directly from the UBC9 or with the added guidance of a SUMO E3 ligase.

Thousands of proteins are SUMOylated and deSUMOylated in a spatially and temporally controlled manner [9–12]. SUMO1 is conjugated to its substrates chiefly as a single conjugate (mono-SUMOylation), with a small proportion in SUMO chains, whereas SUMO2/3 more often form chains (poly-SUMOylation) [13, 14]. Most SUMO1 in mammalian cells appears conjugated to proteins, whereas much of SUMO2/3 is found unconjugated, and their conjugation to substrates is increased following cellular stresses [1, 15]. Some substrates are modified by SUMO1, others by SUMO2/3, and many by both forms [12]. Three broad mechanisms promote differences in substrate-SUMO variant modifications; SUMO-variant-specific SIMs (SUMO interaction motifs) in SUMO E3s and target proteins can bias modification [16–23]; a SUMOylated variant may be protected from isopeptidases after non-specific modification [24, 25]; or SUMO variant and conjugate types may be differentially processed by SUMO proteases (SENPs), some of which exhibit bias.

Successful mitotic signalling is crucial for segregation of the duplicated genome to produce two genetically stable daughter cells, and the SUMO system is critical for successful mitosis. Depletion or inhibition of SAE1/SAE2 or loss, depletion, or mutation of UBC9 results in delays in chromosome alignment, errors in chromosome segregation, and suppression of metaphase to anaphase transitions [26–29] (reviewed in [30]). SUMO1 and SUMO2/3 appear to have differing roles in this portion of the cell cycle. Between 70-600 SUMO2/3 mitotic substrates have been described [31–33]. SUMO2/3 localises to mitotic DNA, and SUMO2/3ylated PARP1 [34] and TOP2 [29] are present on mitotic chromatin. SUMO2/3 is also found at the protein complexes that attach chromosomes to spindle microtubules, the kinetochores, where SUMOylation promotes the recruitment of many proteins, including PLK1 [35], PICH [36] and Aurora B [37, 38]. SUMO1 appears on the structures that help organise microtubules to promote chromosome segregation, the centrosomes, and also along the microtubules of the mitotic spindle [39]. The majority of the observed SUMO1-localisation is thought to be RanGAP1-SUMO1 [39, 40]. Relatively few other mitotic SUMO1 substrates have been reported and include BubR1 [41], Aurora-A [42], PLK1 [35][43] and NuMA [44]. Defective SUMO1ylation of these substrates is associated with prolonged mitosis [43] or spindle abnormalities [40, 44]. How the SUMO conjugation machinery achieves SUMO variant-specific modification of target proteins, particularly in mitosis, is unclear.

Here, we identify a previously undescribed means of SUMO variant-conjugation bias. We find that SAE2 is deacetylated during mitosis and that de-acetylation can be prevented with HDAC6 inhibitor treatment. We find a SAE2 acetyl-analogue, SAE2-K164Q, drives a SUMO2>SUMO1 conjugation bias, resulting in diminished cellular high-molecular weight SUMO1 conjugates in mitotic cells. Cells complemented with SAE2-K164Q exhibit supernumerary structures of the nuclear mitotic apparatus NuMA, multipolar spindles and CENPA-positive micronuclei indicative of poor chromosome segregation. The multipolar spindle phenotype of SAE2-K164Q expressing cells, or of cells treated with HDAC6-inhibitor, can be reversed by SUMO1 overexpression or by expression of a SUMO1-NuMA linear fusion protein, suggesting diminished SUMO1ylation, specifically NuMA SUMO1ylation, is the primary cause of multipolar spindles. These data indicate that SUMO variant bias can be introduced through the SUMO E1, which in turn contributes to mitotic fidelity.

## Results

### SAE1:SAE2-acK164 is deacetylated by HDAC6

Acetylated SAE2-K164 was found in human cervical carcinoma HeLa cells by liquid chromatography mass spectrometry with peptides separated by strong cation exchange chromatography and enriched using a pan-acetyl antibody; acetylated SAE2-K164 was not found following HeLa treatment with 10 Gy ionising radiation [45]. To examine the modification, we generated a monoclonal antibody to the K164-SAE2 peptide, “HP[ac]KPTQRTFPGC”. We tested the antibody on immunoprecipitated (wild-type) WT FLAG-SAE2 and FLAG-SAE2-K164R mutant, noting that the antibody did not detect the mutant (Supp. Fig. 1a). We examined lysates for the acK164-SAE2 signal an hour after irradiation exposure (10 Gy), finding a reduction, with no change detected in total FLAG-SAE2 protein after treatment (Supp. Fig. 1a). Further we noted reduced acK164-SAE2 signal in lysates from cells given the microtubule polymerisation inhibitor nocodazole and harvested by mitotic shake off (Fig. 1a), suggesting de-acetylation is co-incident with early mitosis. We next tested several inhibitors of deacetylases and noted that the inhibitor Panobinostat, a Histone Deacetylase (HDAC) Class I, II, and IV inhibitor, suppressed loss of the acK164-SAE2 signal, as did the selective HDAC6 inhibitor, ACY-738 [46] (Fig. 1a). To test this finding further, we examined whether SAE2 and HDAC6 interact. In immunoprecipitation experiments, endogenous HDAC6 co-immunoprecipitated with FLAG-SAE2 and was enriched further in FLAG-SAE2 precipitates from U2OS cells in mitosis–isolated using nocodazole and mitotic shake-off (Fig. 1b). These data suggest SAE2-K164 de-acetylation occurs through the interaction and activity of HDAC6 and is increased in early mitosis. Next, we addressed whether specific acetylase enzymes contribute to the modification. We tested inhibitors to p300, TIP60, NAT10, and GCN5 and found that no single inhibitor application suppressed the detection of the acK164-SAE2. Combining NAT10, TIP60, and p300 inhibitors reduced the acK164-SAE2 signal, although not completely (Fig. 1c). These data suggest that SAE2-K164 acetylation occurs via the activity of more than one acetylase enzyme.

**Figure 1.**
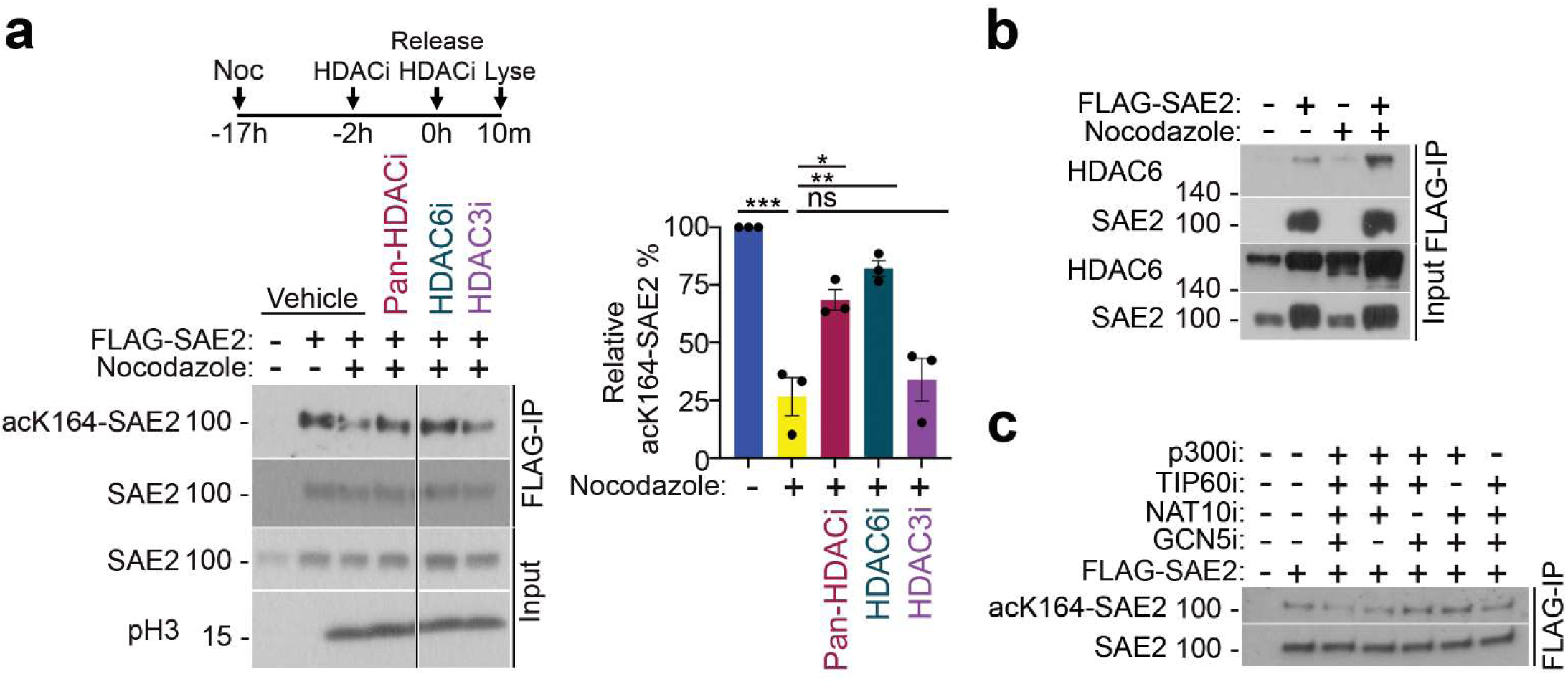
Acetylation of SAE2-K164. **a.** Representative western blot of acetyl-K164-SAE2 (mouse monoclonal) following anti-FLAG immunoprecipitation from U2OS cells and U2OS cells expressing FLAG-SAE2 in unsynchronised cells or cells treated with nocodazole for 17 hours before washing and releasing into mitosis for 10 minutes. HDAC inhibitors, Panobinostat (HDAC class I, II, and IV), ACY-738 (HDAC6), and S7229 (HDAC3), were applied to cells in the last 2 hours of nocodazole synchronisation at 2.5 µM and reapplied upon the release. Antibody specific for phosphorylated histone 3 is used as a marker of mitosis. Quantification of the relative abundance of acetyl-K164-SAE2 relative to the total abundance of SAE2 immunoprecipitated. Error bars = SD; N = 3. *** = p< 0.001, ** = p< 0.01, * = p< 0.05, ns= not significant p>0.05. Statistical significance was calculated using one-way ANOVA. **b.** Western blot analysis of a FLAG-SAE2 co-immunoprecipitation with HDAC6 from U2OS, in the context of an asynchronous cell population or following a 16-hour nocodazole-treatment and mitotic shake-off. **c.** Western blot of acK164-SAE2 following anti-FLAG immunoprecipitation from U2OS cells and U2OS cells expressing FLAG-SAE2 in asynchronous cells treated with indicated combinations of inhibitors against the histone acetyltransferases p300, TIP60, NAT10, and GCN5. Inhibitors were added for 2 hours at 2.5 µM.

### SAE2-K164Q has a SUMO2>SUMO1 bias

Residue K164 of SAE2 forms part of the tunnel through which the C-terminal tails of SUMO proteins access the SAE1:SAE2 catalytic sites in SUMO activation [7, 47, 48] (Supp. Fig. 1b). This proximity led us to query whether K164- SAE2 modification might impact SUMO protein interactions. We generated recombinant (WT) SAE1:SAE2 proteins and a mutant in which the SAE2 element carried a mutation at codon 164 to glutamine (Q) (Supp. Fig. 2a contains SDS-PAGE gels of SEC fractions of these proteins and all the purified proteins generated in the current study). Glutamine resembles the uncharged carbonyl functional group of an acetylated lysine, and although not forming a classical isostere, glutamine has been used as an acetyl-analogue mutant [49–52]). We labelled SAE1:SAE2 and SAE1:SAE2- K164Q with NT-647-NHS for MicroScale Thermophoresis (MST). The technique detects the movement of fluorophore-tagged molecules in a temperature gradient. The thermophoresis of a protein differs from that of its liganded complex, such that MST can be used to quantify interaction dissociation constants [53]. We subjected labelled SAE1:SAE2 to MST analysis with increased concentrations of SUMO1 or SUMO2. Fitting of the data to the change in thermophoresis showed that SAE1:SAE2 has a greater affinity (lower *K_d_*) for SUMO1 (3.7 ± 1.1 µM) than SUMO2 (14.7 ± 1.8 µM; Fig. 2a and Supp. Fig. 3a), consistent with *K_m_* measurements of SUMO1 *Vs* SUMO2 with SAE1:SAE2 [54]. Intriguingly, performing the same analysis with SAE1:SAE2-K164Q revealed a greater affinity (lower *K_d_*) for the SUMO2 (0.4 ± 0.13 µM) and reduced affinity for SUMO1 (28.0 ± 12.19 µM; Fig. 2a and Supp. Fig 3a). These data show codon 164 of SAE2 can influence SAE1:SAE2 affinity for SUMO proteins.

**Figure 2.**
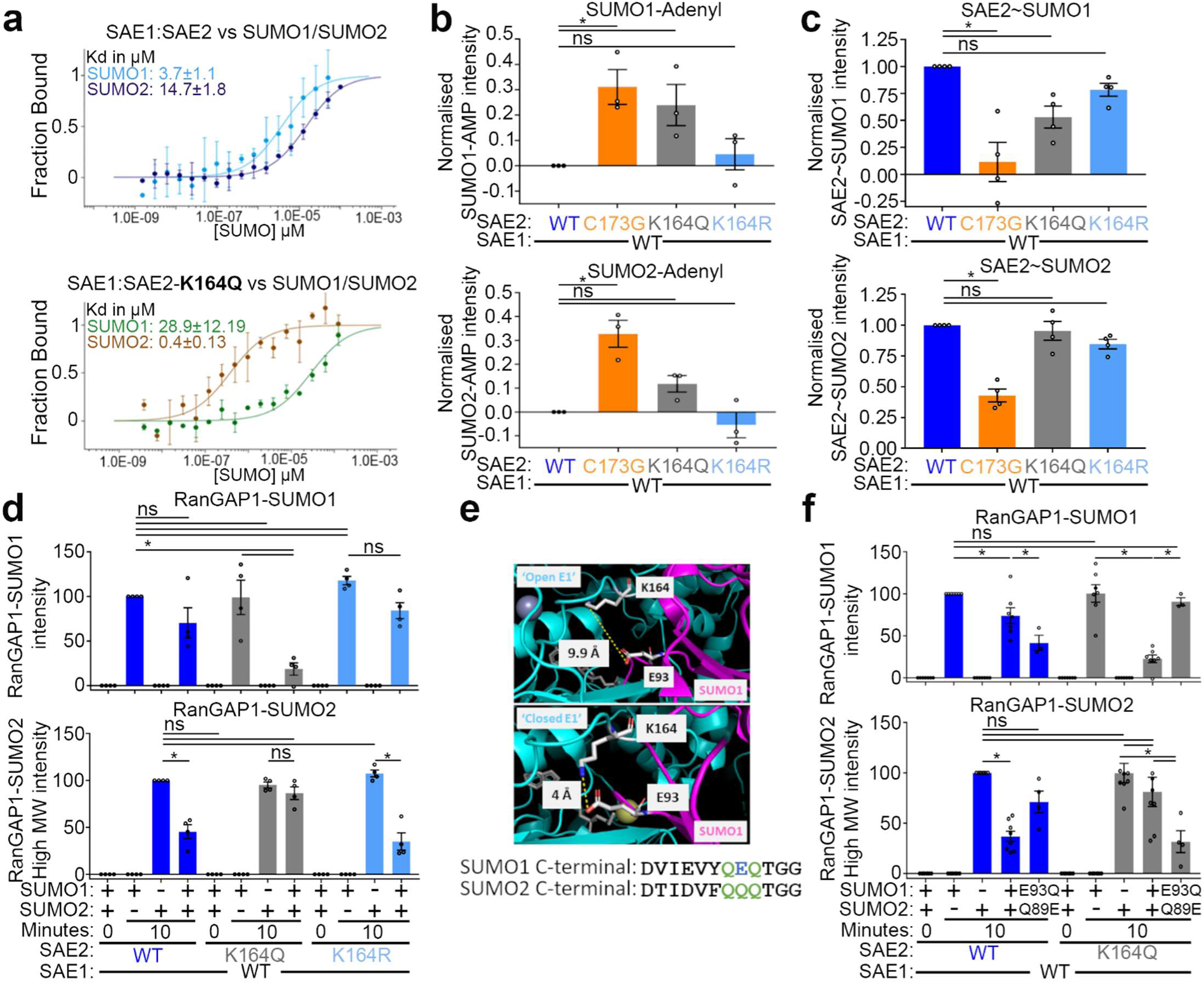
SUMO activation by SAE1:SAE2-K164Q is biased towards SUMO2. **a.** MST analysis of SAE1:SAE2-K164Q affinity with SUMO1 and SUMO2. SUMO1 and SUMO2 concentrations were titrated from 125 µM to 0.00381 µM against 10 nM of NT647-labeled, SAE1:SAE2 (top) or SAE1:SAE2-K164Q (bottom). Plotting of the change in thermophoresis and fitting of the data yielded a *K_d_* in µM shown. N =3, error bars = SD. **b.** *In vitro* SUMO1 adenylation assay, conducted by combining 30 µM SAE1:SAE2-K164 variants with 40 µM SUMO1 or SUMO2, 150 µM BODIPY-ATP, and 1 U pyrophosphatase for 10 minutes at 30°C. Reactions were quenched by the addition of loading buffer and incubation at 95°C for 5 minutes before running samples on 15% SDS-PAGE. Gels were imaged at 488 nm to excite BODIPY-ATP, where a band at 15 kDa was taken to be SUMO-AMP-BODIPY (Supp. Fig. 3b shows a representative gel). N = 3, error bars = SEM, and statistical significance calculated using ANOVA where * = p<0.05. **c.** *In vitro* SUMO loading assays combining 10 μM SUMO with 5 μM SAE1:SAE2-K164 variants or SAE1:SAE2-C173G. Reactions were initiated by the addition of 5 mM ATP at 30°C for 10 minutes and terminated by the addition of reducing agent-free loading buffer and boiling samples. Samples were analysed by SDS-PAGE where fluorescent SUMO bands at 120 kDa were taken to be SAE2∼SUMO, which were quantified and normalised to the WT SAE1:SAE2 condition. **d.** *In vitro* SUMOylation of RanGAP1 (aa398-587) fragment in the presence of 20 μM SUMO1 and/or 20 μM SUMO2, 200 nM SAE1:SAE2, 500 nM UBC9, 10 μM RanGAP1 (aa398-587), and 5 mM ATP, incubated at 30°C for 10 minutes. Supp. Fig. 3c & d show an analysis of band identities in SUMOylation assays. N = 4, error bars = SEM, with statistical significance calculated using one-way ANOVA, * = p<0.05. **e.** Examination of human SUMO E1 K164-SAE2 (Cyan) in ‘open’ conformation (Top) with E93-SUMO1-AMSN (Magenta; PDB ID code 3KYC) and in the ‘closed’ conformation (Bottom) with E93-SUMO1-AVSN (Magenta; PDB ID code 3KYD [48]). In the ‘open’ conformation (Top) the SAE2-K164 ζ-nitrogen and SUMO1-E93 ε-carbonyl are 9.9 Å apart, while the ‘closed’ SUMO E1 conformation shows SAE2-K164 ζ-nitrogen and SUMO1-E93 ε-carbonyl ∼4 Å apart [48], a suitable distance for a noncovalent interaction. Amino acid sequence alignment for SUMO1-E93 and SUMO2-Q89. **f.** *In vitro* SUMOylation RanGAP1 (aa398-587) in the presence of WT SUMO1, and/or SUMO2, SUMO1-E93Q and SUMO2-Q89E (all SUMO variants supplied at 20 μM), 200 nM SAE1:SAE2, 500 nM UBC9, 10 μM RanGAP1 (aa398-587), and 5 mM ATP, incubated at 30°C for 10 minutes. N = 4, error bars = SEM, *p<0.05. Statistical significance was calculated using one-way ANOVA.

To assess the SUMO activation capacity of SAE1:SAE2 *Vs* SAE1:SAE2-K164Q, we generated two additional recombinant SAE1:SAE2 variants, SAE1:SAE2-K164R, and a thiolation catalysis mutant SAE1:SAE2-C173G. We tested SUMO protein adenylation *in vitro* through incubation of SAE1:SAE2 variants with SUMO1 *or* SUMO2 and fluorescently labelled ATP, BODIPY-ATP, followed by the quantification of SUMO1-AMP-BODIPY or SUMO2-AMP-BODIPY [48]. We first tested the SAE1:SAE2-C173G mutant form of the enzyme. The C173G mutant can catalyse SUMO-AMP formation but lacks the catalytic cysteine needed for thioester bond formation and incubation with SAE1:SAE2-C173G resulted in high SUMO1- and SUMO2-AMP-BODIPY levels. In contrast, incubation with WT enzyme produced low levels of SUMO-AMP-BODIPY (Fig. 2b and Supp. Fig. 3b). By comparison, SAE1:SAE2-K164Q showed elevated SUMO1 adenylation, while the levels of SUMO2 adenylation were comparable to those of the WT enzyme (Fig. 2b and Supp. Fig. 3b). Incubation with the structurally similar SAE1:SAE2-K164R mutant resulted in SUMO1-AMP-BODIPY and SUMO2-AMP-BODIPY to a comparable extent observed for WT SAE1:SAE2 reactions.

To test SAE2∼SUMO thioester formation, we combined recombinant SAE1:SAE2-K164 variants with SUMO1-C52A-S9C- Alexa488 or SUMO2-C48A-A2C-Alexa647 [55, 56] in ATP buffer. Using SDS-PAGE under non-reducing conditions, we monitored the formation of a 120 kDa fluorescent SUMO band consistent with loaded SAE2∼SUMO. In these assays, and as expected, SAE1:SAE2-C173G failed to produce a 120 kDa SAE2∼SUMO band (Fig. 2c). Interestingly, the reaction involving SAE1:SAE2-K164Q with SUMO1 showed a reduction in SAE2∼SUMO1 product while the reaction between SAE1:SAE2-K164Q and SUMO2 did not significantly deviate from the WT SAE1:SAE2 reaction (Fig. 2c). These findings imply that the SAE1:SAE2-K164Q mutant can generate adenylated SUMO1, but that it is inefficient at converting SUMO1-AMP to the thioester-linked SAE2∼SUMO1, whereas the ability of SAE1:SAE2-K164Q to process SUMO2 is unaffected.

Next, we wished to test whether SAE2 residue K164 influences substrate SUMOylation, particularly when both SUMO1 and SUMO2 are available. We used SUMO1 and SUMO2, UBC9 and a model substrate, a RanGAP1 fragment (amino acids 398-587). We found all SAE1:SAE2 enzyme variants tested could promote RanGAP1 SUMOylation to a comparable degree after a 10-minute reaction when supplied with single SUMO variants (Fig. 2d). In contrast, when incubated with equimolar amounts of SUMO1 and SUMO2, WT-SAE1:SAE2 or SAE1:SAE2-K164R exhibited a bias to SUMO1 conjugation, whereas the inclusion of SAE1:SAE2-K164Q in the assay resulted in less SUMO1-RanGAP1 modification and more SUMO2-RanGAP1 conjugation (Fig. 2d and Supp. Fig. 3c & d). These data indicate SAE1:SAE2- K164Q drives a SUMO2>SUMO1 conjugation bias.

To address how codon 164 of SAE2 discriminates between SUMO proteins, we examined published structures of SAE1:SAE2-SUMO1-AVSN [47, 48] and noted proximity between lysine 164 of SAE2 and glutamate 93 of SUMO1 (Fig. 2e). Intriguingly, the equivalent residue in SUMO2 is glutamine (Fig. 2e). To test the hypothesis that SAE2 codon 164 discriminates SUMO proteins through these residues we swapped them, creating SUMO1-E93Q and SUMO2-Q89E. Remarkably, the inclusion of these substitutions switched the conjugation bias of SAE1:SAE2-K164Q from SUMO2 to SUMO1, and also reversed the SUMO1 bias of WT SAE1:SAE2 from SUMO1 to SUMO2 (Fig. 2f). These data are consistent with the idea that codon 164 of SAE2 contributes discriminative interactions with SUMO protein C-terminal regions to influence SUMO protein variant conjugation.

### SAE1:SAE2-K164 supports mitotic fidelity

To address the potential cellular consequences of K164 modification, or lack of modification, we generated a dual expression system for siRNA-resistant SAE1:SAE2. We made WT E1, the acetyl-analogue SAE1:SAE2-K164Q, the thiolation-inactive SAE1:SAE2-C173G [4] and SAE1:SAE2-K164R (Supp. Fig 4a). The latter was generated to maintain structural similarity and test the degree to which lysine 164 modification, which will be lost on both K164Q and K164R mutations, is needed. We then tested for potential defects in several cellular phenotypes driven by SUMO modifications. We first examined the survival of cells exposed to heat shock, where SUMOylation of hundreds of proteins is required to survive the stress [57]. Depletion of SAE1:SAE2 resulted in cellular sensitivity to 43°C exposure that could be suppressed by complementation with SAE1:SAE2, SAE1:SAE2-K164R, or SAE1:SAE2-K164Q, but not SAE1:SAE2-C173G (Supp. Fig. 4b), indicating little or no influence of the SAE2-K164, or K164 modification on the heat shock response. It may be relevant that the heat shock response involves the conjugation of SUMO2/3 over SUMO1 [58][57].

The cellular response to DNA double-strand breaks requires many SUMOylation events [3]. As expected, cells exposed to irradiation showed delayed resolution of the DNA damage marker γH2AX on depletion of UBC9 or the SUMO E3 ligase PIAS1 (Supp. Fig. 4c). Intriguingly, however, we found no impact of SAE1:SAE2 depletion or SAE2- complementation, even with SAE1:SAE2-C173G complementation, on γH2AX kinetics (Supp. Fig 4c). Consistent with these observations, we observed only a slight impact of SAE1:SAE2 depletion on the repair of enzymatically generated DNA double-strand breaks (Supp. Fig. 4d). In the light of the requirement for SUMOylation in the DNA damage response, these results are surprising, nevertheless, they are consistent with the report that SUMO E1 inhibition has no impact on the cellular response to DNA damaging agents, cisplatin or hydroxyurea [26].

We next considered the potential influence on mitosis since the HDAC6:SAE2 interaction and acK164-SAE2 de-acetylation were co-incident with early mitosis (Fig. 1a & b). To examine the impact of SAE1:SAE2 on mitosis more precisely, we generated a second complementation system to allow acute E1 inhibition. In this system, U2OS cells express a SAE2 mutant, SAE2-S95N-M97T, that we name “FLAG-SAE2r”, which provides resistance to the SUMO E1 inhibitor, ML792 [26]. Into FLAG-SAE2r, we added SAE2-C173G, SAE2-K164Q and SAE2-K164R mutations. We assessed the impact of SAE2-K164 mutations on mitotic SUMOylation, examining cells bearing FLAG-SAE2r variants synchronised in prometaphase with nocodazole for 16 hours with the concurrent addition of ML792, examining cells 10 minutes after release from nocodazole treatment (Fig. 3a). In all FLAG-SAE2r-complemented cells, high-molecular-weight immunoprecipitated SUMO2 conjugates were equivalent despite the lower expression of FLAG-SAE2r-K164R and FLAG-SAE2r-K164Q mutants (Fig. 3a, right-hand panel). FLAG-SAE2r complemented cells showed high-molecular weight immunoprecipitated SUMO1 conjugates, whereas the lesser expressed FLAG-SAE2r-K164R exhibited less high-molecular weight SUMO1 (Fig 3a, left-hand panel). Despite the equivalent expression levels of FLAG-SAE2r-K164Q and FLAG-SAE2r-K164R, high-molecular-weight SUMO1 was almost undetectable in the precipitates from cells complemented with FLAG-SAE2r-K164Q. These data indicate that SUMO1 conjugate generation is sensitive to E1 levels and to SAE2-K164Q.

**Figure 3.**
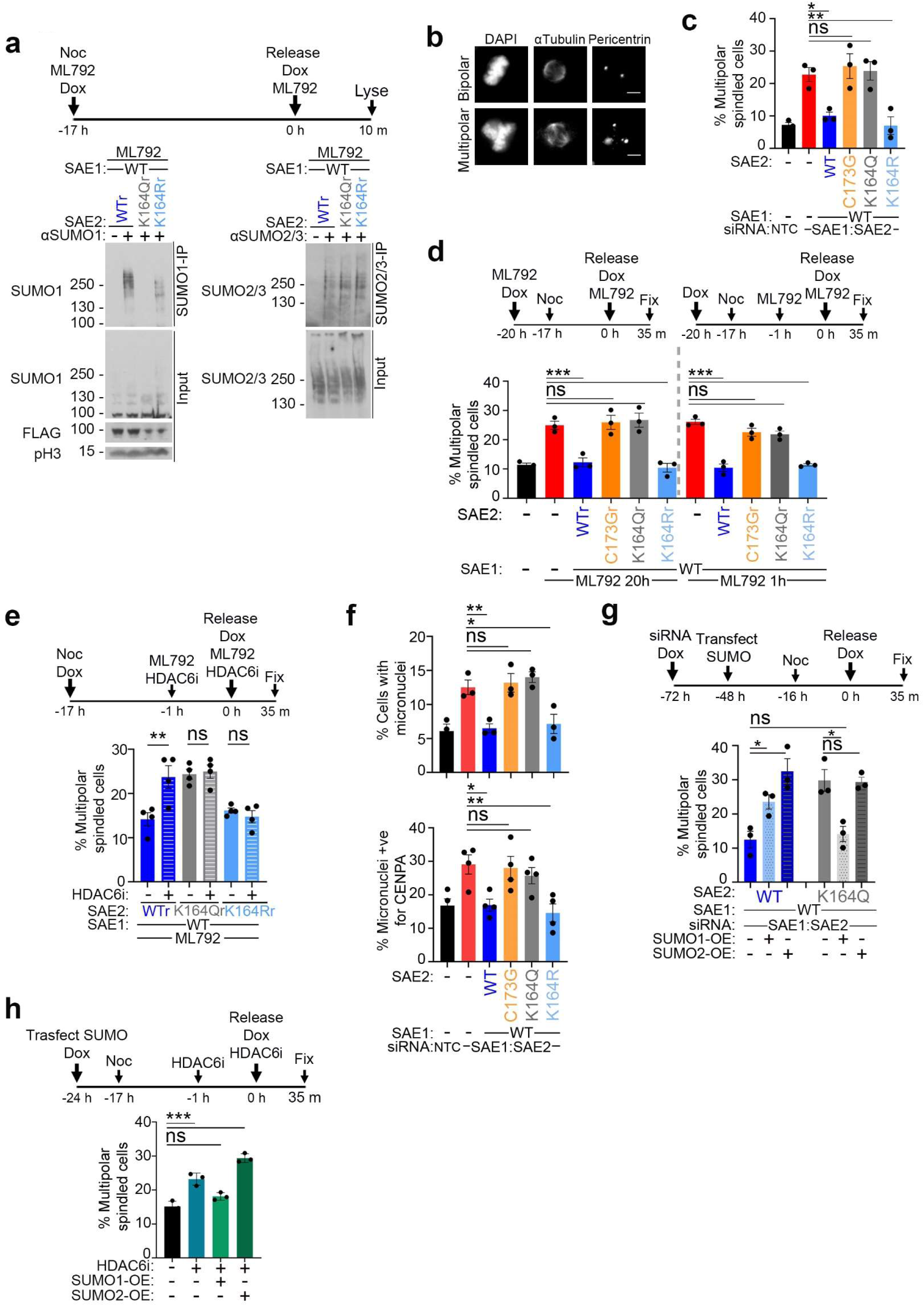
K164-SAE2 directs mitotic fidelity. **a.** Immunoprecipitation of endogenous mitotic SUMO conjugates in cells treated with ML792 and expressing Flag-SAE2 constructs resistant to the inhibitor. ML792 resistance is denoted by (r). The presence of Flag-SAE2r is represented by (WTr), Flag-SAE2r-K164Q by KQr, and Flag-SAE2r-K164R by KRr. Diagram, top, illustrates the timing of inhibitor and induction agent addition. **b.** Representative images of immunofluorescent analysis of mitotic spindle formation. Five-micron scale bar is shown as a white line. **c.** The percent of laterally presented metaphase and anaphase cells exhibiting multipolar spindles in cells complemented with SAE2 variants in RNAi-treated cells, as shown. Error bars SEM; N = 3. Significance calculated using one-way ANOVA, *= p≤ 0.05, **= p≤ 0.01, ns = not significant > 0.05. Data from 3 independent biological repeats. n>50 cells per condition analysed from a minimum of four fields of view per biological repeat. **d.** The percent of laterally presented metaphase and anaphase cells exhibiting multipolar spindles. Cells exposed to different durations of ML792 prior to release from nocodazole: long treatment totalling 20h (added 4h before nocodazole) is displayed to the left-hand side, short treatment totalling 1h (added during the last hour of nocodazole) presented to the right-hand side. ML792 replaced onto cells after release from nocodazole. Error bars SEM; N = 3. Significance calculated using one-way ANOVA, ***= p≤ 0.001, ns = not significant > 0.05. Data from 3 independent biological repeats. n>50 cells per condition from a minimum of four fields of view per biological repeat. **e.** The percent of laterally presented metaphase and anaphase cells exhibiting multipolar spindles in cells treated with ML792 and complemented with WTr, Flag-SAE2r-K164Q, and Flag-SAE2r-K164R with and without the addition of HDAC6 inhibitor. Error bars SEM; N = 4. Significance calculated using one-way ANOVA, **= p≤ 0.01, ns = not significant > 0.05. Data from 4 independent biological repeats. n>50 cells per condition from a minimum of four fields of view per biological repeat. **f.** Analysis of micronuclei in asynchronous siRNA-resistant SAE2 variant cells. The percentage of total cells with one or more micronuclei is plotted (Top). Error bars SEM; N=3. Significance calculated using one-way ANOVA, *=p≤ 0.05, **≤ 0.01, ns = not significant > 0.05. Data from 3 independent biological repeats. Cells from at least 4 fields of view were analysed. n>400 total cells per condition per biological repeat. The percentage of micronuclei positive for CENPA in asynchronous siRNA-resistant SAE2 variant cells (Bottom). Error bars SEM; Significance calculated using one-way ANOVA, N=4 *=p≤ 0.05, **≤ 0.01, ns = not significant > 0.05. Data from 4 independent biological repeats. n>50 micronuclei per condition per biological repeat. **g.** The percent of metaphase cells with multipolar spindles in cells expressing siRNA-resistant SAE2 variants after release from nocodazole, with or without SUMO1 or SUMO2 overexpression. Timeline of the experiment depicted above. Error bars SEM; N=3 *=p≤ 0.05, **≤ 0.01, ns = not significant > 0.05. Data from 3 independent biological repeats. n>50 cells per condition from at least four fields of view per biological repeat. **h.** Average percent of metaphase cells with multipolar spindles in cells treated with HDAC6 inhibitor, with or without 24 hr SUMO1 or SUMO2 overexpression. Timings of the experiment are displayed above. HDAC6 inhibitor added 1 hr prior nocodazole and replaced onto cells for the duration of mitotic release Error bars SEM; Significance calculated using one-way ANOVA, *=p≤ 0.05, **≤ 0.01, ns = not significant > 0.05. Data from 3 independent biological repeats. n>50 cells per condition from at least four fields of view per biological repeat.

SUMO conjugation promotes correct spindle assembly and chromosome segregation [30, 59]. To assess if SAE2-K164 impacts mitotic spindle assembly, we first depleted endogenous SAE1:SAE2 and expressed siRNA-resistant SAE1:SAE2 variants before synchronisation with nocodazole, wash-out and immunostaining for components for the mitotic spindle machinery, α-tubulin and pericentrin, inspecting cells in metaphase and anaphase. Depletion of SAE1:SAE2 increased the proportion of cells with multipolar spindles, which were suppressed by the expression of SAE1:SAE2 or SAE1:SAE2-K164R (Fig. 3b & c). However, neither SAE1:SAE2-C173G nor SAE1:SAE2-K164Q expression suppressed their formation (Fig. 3c). These data suggest that SAE2-K164Q is less able to support bipolar spindle formation, whereas SAE2-K164R, promotes normal spindle polarity.

To begin to address how spindle defects might arise, we asked when E1 activity is required. Multipolar spindle formation can be driven by centrosome amplification in S-phase or G2 and also by aberrant initiation of spindle assembly occurring on or after nuclear envelope breakdown [60]. We tested FLAG-SAE2r expressing cells and added the E1 inhibitor in the last hour of nocodazole treatment, before wash-out, as well as before nocodazole exposure (Fig. 3d). Under both conditions, FLAG-SAE2r-WT and FLAG-SAE2r-K164R, but not FLAG-SAE2r-C173G or FLAG-SAE2r-K164Q, suppressed the formation of multipolar spindles (Fig. 3d). These data discount defects in S-phase or G2 as the drivers of multipolar spindle generation. Instead, they suggest that absent SUMOylation and the impact of SAE2-K164Q disrupt a process needed directly before, or during re-polymerisation of the microtubule network.

Next, we tested the role of HDAC6 de-acetylation activity in the formation of multipolar spindles in the context of FLAG-SAE2r variants. Cells containing FLAG-SAE2r variants were synchronised with nocodazole for 16 hours and treated for the last 1 hour with ML792 with or without HDAC6 inhibitor, ACY-738, before releasing into mitosis (Fig. 3e). HDAC6 inhibition resulted in a significant increase in multipolar spindles in the FLAG-SAE2r-WT cells, consistent with previous reports of HDAC6 inhibition [61]. However, importantly, HDAC6 inhibitor induced no significant change to multipolar spindle levels in cells expressing either K164 mutant. In FLAG-SAE2r-K164Q expressing cells, the high levels of multipolar spindles remained high, whereas in cells expressing FLAG-SAE2r-K164R, multipolar spindles remained low despite HDAC6 inhibitor treatment (Fig. 3e). Thus, SAE2-K164 status imparts insensitivity of spindle assemblies to HDAC6 inhibition, suggesting de-acetylation of SAE2-K164 acts to promote bipolar spindles.

Cells with more than two spindle poles may segregate chromosomes poorly, and misaligned, lagging, or bridge chromosomes can become isolated as encapsulated DNA fragments, known as micronuclei [62]. We examined SAE1:SAE2 siRNA complemented cells for micronuclei and found that SAE1:SAE2 and SAE1:SAE2-K164R complementation, but not SAE1:SAE2-C173G or SAE1:SAE2-K164Q, suppressed increased numbers of micronuclei (Fig. 3f, top panel), consistent with the notion that inappropriate spindle formation results in micronuclei in these cells. To gain further insight into the contents of the micronuclei, we stained for the centrosome component, CENPA, and the DNA-damage marker γH2AX. We observed no enrichment for γH2AX-containing micronuclei in SAE1:SAE2 depleted or complemented cells (Supp. Fig. 4e), suggesting no increased chromosome fragments in the micronuclei observed. However, approximately 1/3^rd^ of micronuclei in SAE1:SAE2 depleted or SAE1:SAE2-C173G or SAE1:SAE2-K164Q complemented cells showed staining for CENPA (Fig. 3f, bottom panel), suggesting a proportion of the micronuclei contain centric chromosomes.

Given the role of K164 in SAE2 promoting SUMO1ylation in the context of SUMO1:SUMO2 competition (Fig. 2f), we considered whether the defects of cells complemented with SAE2-K164Q, which show reduced SUMO1 conjugates might be rescued by changing the balance between SUMO variant expression. Remarkably, SUMO1, but not SUMO2, overexpression suppressed the formation of multipolar spindles in SAE1:SAE2-K164Q complemented cells (Fig. 3g), consistent with the notion that SUMO1ylation driven by SAE2-K164 is critical to spindle regulation. Intriguingly, both SUMO1, and in particular, SUMO2 over-expression also increased the number of multipolar spindles in cells complemented with WT SAE1:SAE2, suggesting that disrupting SUMO variant balance disturbs normal spindle assemblies. Consistent with the idea that HDAC6 drives SAE2 de-acetylation to support mitotic SUMO1ylation, SUMO1 overexpression suppressed multipolar spindle formation in cells treated with HDAC6 inhibitor (Fig. 3h).

### SUMO1-NuMA fusion suppresses spindle defects in HDCA6-inhibitor-treated and SAE2-K164Q-complemented cells

Our findings of a SAE2-K164:SUMO1 dependency in mitotic spindle organisation led us to consider whether known specific SUMO1ylated substrate(s) may explain the defect in HDAC6 suppressed or SAE1:SAE2-K164Q-complemented cells. Among the known SUMO1 conjugates in mitosis, the RanBP2/RanGAP1-SUMO1/UBC9 complex and NuMA have previously been associated with the promotion of bipolar mitotic spindles [44, 63]. The remainder of the known SUMO1 conjugates in mitosis, BubR1, Aurora-A and PLK1 promote mitotic timing or microtubule polymerisation [41–43]. RanGAP1-SUMO1 is stable over several cell cycles upon treatment with ML792 [26]) or siUBC9 [64]. Thus, we reasoned that downregulated RanGAP1-SUMO1 is not likely to be responsible for the mitotic defects observed following acute E1 suppression. In contrast, on inspection of the nuclear mitotic apparatus (NuMA) protein in mitotic cells we observed a prominent band at the expected molecular weight for unmodified NuMA (∼238 kDa) and a band at ∼250 kDa that was lost on treatment with the E1 inhibitor ML792 (Fig. 4a), consistent with the previously noted mitotic NuMA SUMO1ylation [44]. NuMA aids the clustering of the microtubule fibre minus ends at the spindle poles around the centromere in early mitosis, and NuMA dysfunction causes spindle pole-focusing defects, lagging chromosomes in anaphase and micronuclei formation [65–67]. Further, a SUMOylation-deficient NuMA mutant is defective in recruitment to spindle poles, in microtubule bundling, and multiple spindles are induced during mitosis [44].

**Figure 4.**
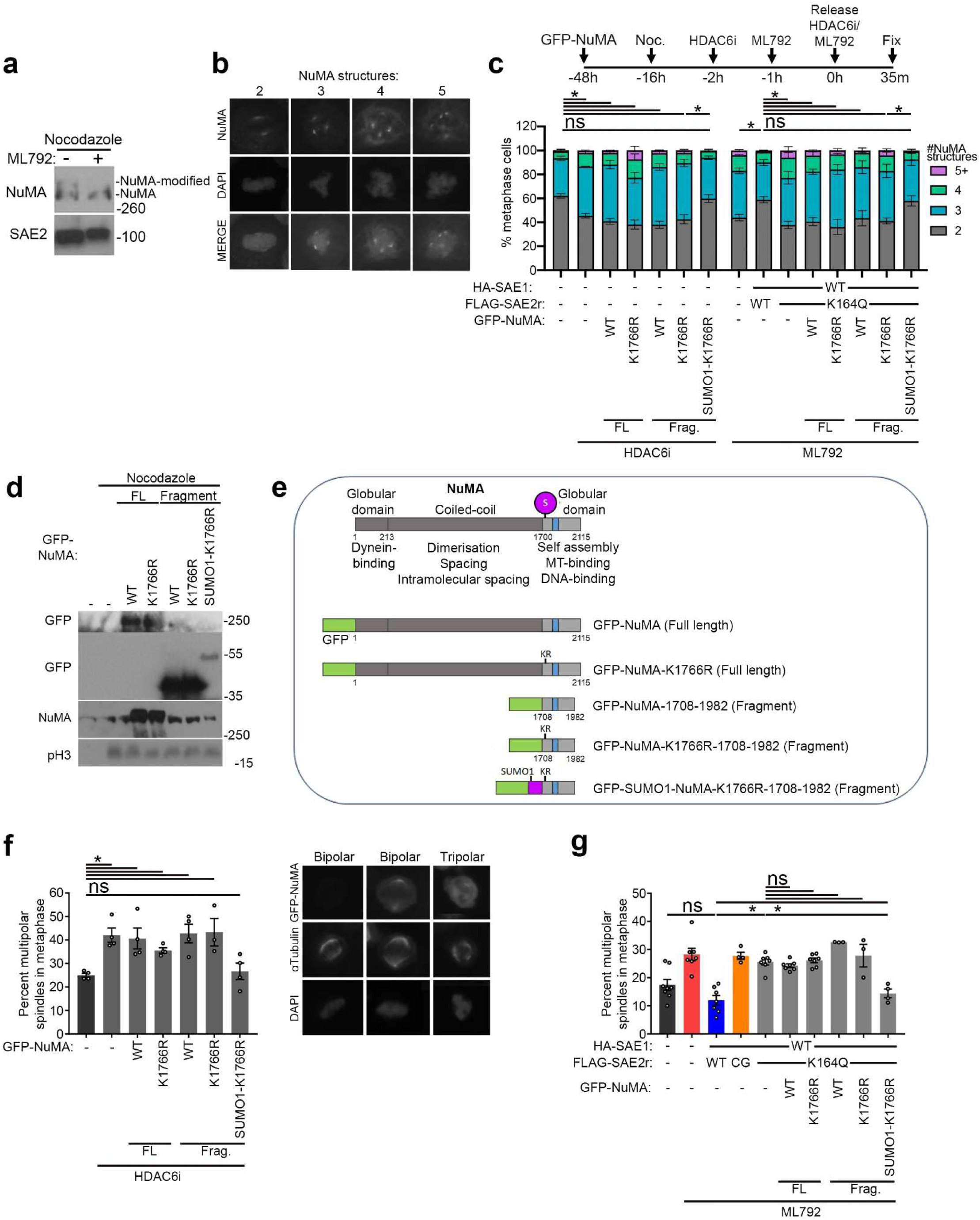
Mitotic defects incurred by SAE2-K164Q of HDAC6 inhibition are corrected by GFP-SUMO1-NuMA fragment expression. **a.** Western blot analysis of U2OS treated with nocodazole ±5 µM ML792 for 16 hours. Mitotic cells were harvested by mitotic shake-off and lysed in loading buffer, and western probed for NuMA and SAE2. **b.** Images show examples 2, 3, 4, and 5 NuMA structures in metaphase cells immunostained for NuMA and stained with Hoechst. **c.** Graph left shows the average percentage of the metaphase cell population with 2 (grey), 3 (blue), 4 (green), and 5+ (purple) NuMA structures in untransfected cells treated with 2.5 μM HDAC6i or cells expressing FLAG-SAE2r-K164Q and treated with 1 μM ML792, with full length or C-terminal fragments of GFP-NuMA, GFP-NuMA-K1766R, and GFP-SUMO1-NuMA. Mean plotted with error bars as SEM for N= 4 independent repeats (>50 cells counted per condition) with one-way ANOVA used to assess the statistical significance for the percentage of metaphase cells with 2 NuMA structures where * = p< 0.05 and ns = p>0.05. **d.** Western blot for the GFP-tag showing U2OS transfected with GFP-NuMA constructs enriched in mitosis with nocodazole. The GFP-NuMA and GFP-NuMA-K1766R proteins are full-length, and the C-terminal 1708-1982 fragments are also shown, as well as the GFP-SUMO1-NuMA-K1766R fragment. **e.** Diagram of NuMA monomer indicating the dominant SUMO1ylation site (purple) at K1766 in the C-terminal ‘self-assembly’ domain as identified by Seo *et al*. (2014). a residue in the C-terminal ‘oligomerisation/self-assembly’ domain and adjacent to the ‘clustering motif’, NuMA-E1768-I1778. Shown are the GFP-NuMA constructs, including full-length (FL) GFP-NuMA, GFP-NuMA-K1766R, as well as C-terminal fragments (Frag.) of NuMA, GFP-NuMA-1708-1982, GFP-NuMA-K1766R-1708-1982, and GFP-SUMO1-NuMA-K1766R-1708-1982 linear fusion. **f.** Graph left shows the average percentage of the metaphase cell population with multipolar spindles as assessed by α-tubulin structures in cells treated with 2.5 μM HDAC6 inhibitor, ACY-738, transfected with full-length GFP-NuMA, GFP-C-terminal fragments or GFP-SUMO1-NuMA-K1766R fusion construct. N = 4 independent experiments, bars = SEM., and statistical significance calculated using one-way ANOVA where * = p< 0.05 and ns = p>0.05. Images right show representative metaphase cells expressing WT GFP-NuMA, immunostained for αtubulin, showing bipolar and multipolar (3) spindles. **g.** Mean percentage of the metaphase cell population with multipolar spindles in U2OS expressing FLAG-SAE2r-K164Q treated with 1 μM ML792 with GFP full-length NuMA, C-terminal fragment NuMA variants or SUMO1 fused C-terminal NuMA fragment and stained for αtubulin. N = 4, bars = SEM, and statistical significance was calculated using one-way ANOVA where * = p< 0.05 and ns = p>0.05.

We first considered NuMA assemblies (Fig. 4b) since >2 indicates disordered spindle organisation [66, 67]. We inspected NuMA in HDAC6-inhibitor and ML972-treated and complemented metaphase cells. HDAC6-inhibitor treatment reduced the proportion of cells bearing 2 NuMA structures whilst increasing the incidence >2 assemblies (Fig. 4c, left-hand side). This observation was also seen in cells treated with ML792 (Fig. 4c, right-hand side). Importantly, complementation with FLAG-SAE2r-WT but not FLAG-SAE2r-K164Q was able to restore the percentage of cells with 2 NuMA structures to untreated levels and reduced the number of cells with >2 NuMA structures (Fig. 4c).

To test if SUMOylation of NuMA relates to NuMA structure and spindle defects in HDAC6i-treated cells and cells complemented with FLAG-SAE2r-K164Q, we expressed a series of NuMA constructs. The major SUMO1ylation site on NuMA is K1766 [44], so we expressed GFP-tagged full-length NuMA and full-length GFP-NuMA-K1766R. Then, since many of the functional roles of the protein are encoded in the globular C-terminus, and a C-terminal fragment can perform the mitotic roles of full-length NuMA [44], we also expressed a GFP-tagged NuMA C-terminus (amino acids 1708-1982). We tested this GFP-C-terminal NuMA fragment either as a WT protein, bearing the SUMO conjugation site, K1766R, mutation, or bearing the K1766R mutant and carrying SUMO1 fused between the GFP and NuMA fragment, to mimic K1766-SUMO1ylated-NuMA (Fig. 4d & e), as previously described [44].

Expression of WT GFP-NuMA, GFP-NuMA-K1766R or the WT or K1766R mutant C-terminus had little impact on NuMA structure numbers in HDAC6 inhibitor-treated cells or in ML792-treated cells complemented with FLAG-SAE2r-K164Q, where cells with >2 structures remained high (Fig. 4c-e). Remarkably however, expression of the NuMA C-terminal fragment bearing the SUMO1 fusion, GFP-SUMO1-NuMA-K1766R, suppressed NuMA structures in both contexts, resulting the majority of HDAC6-inhibitor and the majority of FLAG-SAE2r-K164Q complemented cells exhibiting just 2 NuMA structures (Fig 4c-e). Thus, the expression of a SUMO1-NuMA fusion can prevent both the harmful impact of HDAC6 inhibition and complementation with SAE2-K164Q, to support bipolar NuMA structures.

Finally, we tested the GFP-NuMA constructs for their ability to suppress multipolar spindles in HDAC6 inhibitor-treated or FLAG-SAE2r-K164Q complemented cells. As for NuMA structures, we found bipolar spindles were restored and multipolar spindles reduced in cells expressing the GFP-SUMO1-NuMA-K1766R fragment (Fig. 4f & g). Thus, the SUMO1-NuMA fusion can also improve bipolar spindle formation in conditions of HDAC6 inhibition or FLAG-SAE2r- K164Q complementation.

## Discussion

The SUMO E1 enzyme differs from most other ubiquitin-like modifier-activating enzymes in that it activates related but different modifiers. Here, we find a mechanism describing how the SUMO E1 can direct SUMO protein bias. In our *in vitro* assays, an acetylation-mimic, SAE1:SAE2-K164Q, displays accumulation of SUMO1-adenyl, a reduced ability to form the SAE2∼SUMO1 thioester and biased RanGAP1-SUMO2ylation in reactions containing SAE1:SAE2-K164Q and SUMO1 and SUMO2. We find that the E1 enzyme can discriminate between SUMO variants when bearing SAE2-K164Q through the SUMO C-terminal tail residues, SUMO1-E93 and SUMO2-Q89. Our findings are similar to the discrimination that the NEDD8 E1(APPBP1–UBA3) employs to maintain the specificity of NEDD8 over the 55% identical ubiquitin. APPBP1–UBA3 selectively binds NEDD8-A72 and not ubiquitin-R72 [69]. Walden *et al.* (2003) speculated that other E1 enzymes may employ similar modifier discrimination [69], and indeed NEDD8-A72 and Ub-R72 residues align with SUMO1-E93 and SUMO2-Q89 (Supp. Fig. 5a).

In structural assessments of the E1 with SUMO, the E1 assumes an adenylation catalysing ‘open’ conformation and a thioester bond catalysing ‘closed’ conformation [7, 48]. The open conformation suggests hydrogen bonding between SAE2-R119-Y159 and SUMO1-E93 [7], with the closed exhibiting a closer association, and potential hydrogen bonding of SAE2-K164 and SUMO1-E93/SUMO2-Q89 [47, 48] (Supp. Fig. 5b), this is consistent with our data showing SAE1:SAE2-K164Q adenylates SUMO1 and squanders SUMO1-Adenyl before SAE2∼SUMO1 thioester formation. However, as SAE2-K164Q also influences SUMO variant affinity in favour of SUMO2 in the absence of ATP, the SAE1:SAE2 confirmation before SUMO adenylation also appears to contribute some bias. We speculate that both the open and closed SUMO E1 conformations exist in solution.

Our data do not support a positive role for K164 modification, whether acetylated or carrying another modification, since K164R mutant SAE2 has little phenotype in our hands. In most tissues, free SUMO2/3 pools are readily available [9], so it is unclear whether increasing affinity for SUMO2/3 of itself has any biological function. Nevertheless, K164 is highly conserved (e.g. in *S. cerevisiae*, *D. melanogaster*, *D. rerio* and *H. sapiens*), so we do not discount that acetylation of K164 is employed to favour SUMO2/3ylation following another cellular stress. We also do not rule out other regulatory modifications of the site, since modification with ubiquitin/SUMO would be expected to have an inhibitory impact on activity.

We show that the SUMO E1 enzyme is deacetylated at K164-SAE2 after ionising radiation and in cells synchronised in early mitosis. While we find a little role for the E1 in double-strand DNA break repair or in modulating markers of DNA damage after irradiation, we see a clear association with mitotic fidelity. De-acetylation of acK164-SAE2 can be suppressed by HDAC6 inhibition and our findings strongly align HDAC6 activity with SAE2-K164, since mutation of K164 can overcome the impact of HDAC6 inhibition. Intriguingly, HDAC6 is one of the few histone deacetylases found primarily in the cytoplasm, where it catalyses the removal of acetyl groups from substrates, including α-tubulin and HSP90 [70, 71]. As the SUMO E1 enzyme is predominately nuclear [72], we speculate that the interaction between acK164-SAE2 and HDAC6 is increased by nuclear envelope breakdown at the end of prophase in mitosis.

High-molecular weight SUMO1ylated substrates are severely suppressed in mitosis compared to interphase [39], and we required enrichment of SUMO1 to observe high-molecular weight conjugates by immunoblot. We found that the relatively low-levels of SUMO1-conjugates in mitosis are decreased yet further by complementation with the SAE2- K164Q mutant, suggesting that E1 de-acetylation helps support what little SUMO1ylation there is. Why SUMO1 modification is so severely downregulated in mitosis is not clear; mechanisms related to changes in PIAS family activity, protease activity or the dilution of SUMO1 availability on loss of the nuclear membrane are possible.

SAE2-K164 suppression of abnormal multipolar spindles and CENPA-positive micronuclei aligns with previous findings that SUMO E1 depletion or inhibition drives mitotic defects [26, 27]. We find SUMO1 overexpression suppresses multipolar spindles in SAE2-K164Q complemented cells and in HDCA6-inhibitor treated cells, suggesting SUMO1ylation is the preferred variant that is not acutely compensated for by SUMO2/3. Intriguingly in WT cells, expression of SUMO proteins, particularly SUMO2, also results in multipolar spindles, suggesting disruption of variant balance, particularly SUMO2>SUMO1 can also disrupt bipolar spindle development.

We observe that E1 suppression drives increased NuMA structures. NuMA has both mitotic and interphase roles. In interphase, a form of NuMA missing one of its C-terminal microtubule-binding regions can nevertheless contribute to single-stranded DNA break repair [73]. In contrast, a NuMA C-terminal fragment is sufficient to establish bipolar spindles in metaphase cells [44]. The major NuMA SUMO1ylation site is at K1766 (within P-K-V-E = in a SUMO conjugation consensus site, ψ-K-x-E) [44], overlapping with NuMA’s clustering motif, E1768-P1778 [67, 74]). We see that both the defect of extra NuMA structures and multipolar spindles in conditions of HDAC6 inhibition or SAE2- K164Q complementation are suppressed by the presence of the linear GFP-SUMO1-NuMA C-terminal fragment fusion. Our findings thus provide a mechanistic explanation for the previous observation of NuMA-dependent multipolar spindles induced by HDAC6 inhibition [61]. While we do not discount the requirement for other acutely SUMO1ylated mitotic substrates in SAE2-K164Q complemented cells or HDAC6-inhibited cells, our data suggest SUMO1-NuMA can accomplish much of the role(s) that any other SUMO1-substrate(s) might perform in supporting bipolar mitotic spindle assembly. How NUMA SUMO1ylation specifically supports spindle assembly is unexplored. It may be relevant that NuMA contains a consensus SIM motif, ‘IINI’, (residues 1814-1817), that may encourage SUMO-SIM self-assembly and, we speculate, promote the phase-separation that concentrates proteins for mitotic spindle assembly [75].

The SUMO E1 inhibitor TAK-981 (Subasumstat) is a first-in-class SUMOylation inhibitor [76]. It is currently under clinical trials for solid tumours and is an exciting prospect for cancer treatment, particularly when coupled with immune-checkpoint inhibitors [77]. Our findings suggest that modification of the E1 can influence its ability to bias SUMO variant usage and have led us to a model in which SAE2-K164 de-acetylation by HDAC6 in mitosis improves SUMO1ylation, influencing the crucial substrate NuMA, where its SUMO1ylation acts to promote clustering to establish bipolar mitotic spindles (Fig 5). These findings offer a framework to investigate whether the HDAC6-E1- SUMO1-NuMA pathway is part of the efficacy or adverse effects [78] associated with TAK-981 treatment.

**Figure 5.**
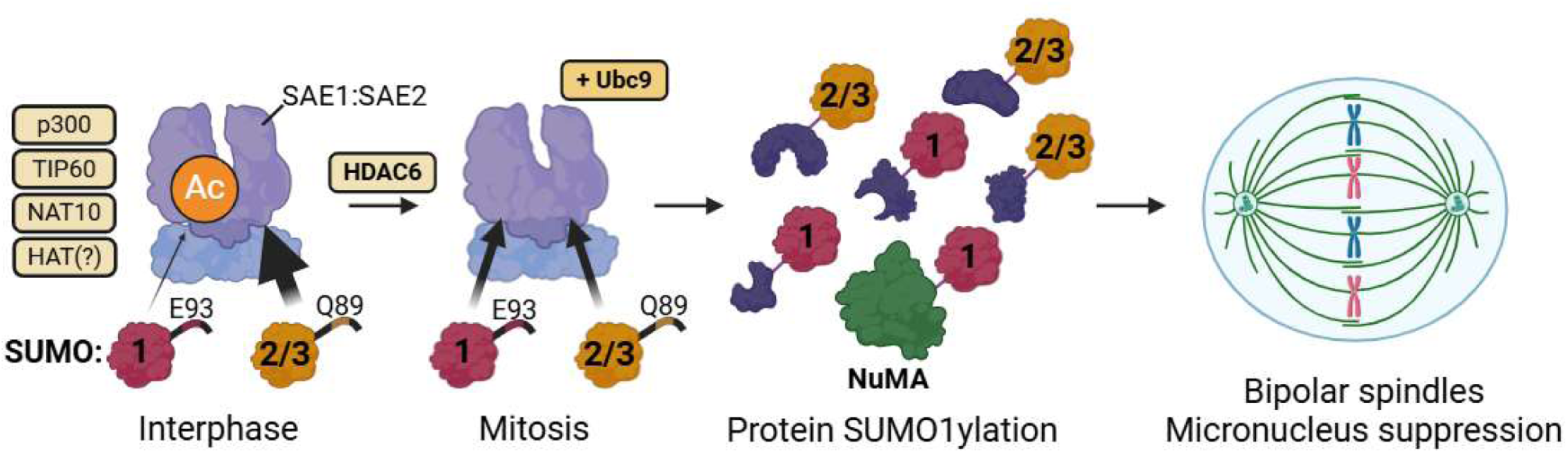
SAE1:SAE2-acK164 is downregulated in mitosis to promote mitotic fidelity. SAE1:SAE2-acK164 is present in mammalian cells due to histone-acetyltransferase activities including p300, TIP60, and NAT10. SAE1:SAE2-acK164 biases the activation of SUMO2 over SUMO1. During mitosis, SAE1:SAE2-acK164 is downregulated in an HDAC6- dependent manner to improve activation of SUMO1 and enable protein SUMO1ylation including NuMA-SUMO1 formation, to promote bipolar mitotic spindles and genomic stability. Created with Biorender.

## Supporting information

Licence for biorender image

## Acknowledgements

Grant funding. Wellcome Trust 206343/Z/17/Z (M.J, A.W, A.J.G), University of Birmingham (A.L). BBSRC: BB/V01983X/1, BB/S017283/1 BB/P009840/1 (TJK). Midlands Integrative Biosciences Training Partnership - BB/M01116X/1 (P.W. & B.F.C). We thank Jeremy Stark (City of Hope, Duarte U.S.A.) for U2OS DR-GFP and NHEJ-EJ5 cells and Cheol Yong Choi (Sungkyunkwan University, Republic of Korea) for pC3-GFP-NuMA constructs. We thank Helen Walden (University of Glasgow, U.K.) and Chris Lima (Sloan Kettering Institute, U.S.A) for their invaluable discussions about the project. We also thank the Microscopy and Imaging Services at Birmingham University (MISBU) and the UoB Flow Cytometry Services (UoBFC) in the Tech Hub facility for microscope and FACS support and maintenance. Model schematic in Fig. 5 created in BioRender. Lanz, A. (2025) https://BioRender.com/s07y449.

## Author contributions

A.K.W. (Immunofluorescent microscopy and analysis, DR-GFP and NHEJ-EJ5 reporter assays, immunoprecipitations, western blotting, site-directed mutagenesis, generation of stable cell lines); A.J.L (Site-directed mutagenesis, generation of stable cell lines, acK164-SAE2 antibody development, immunoprecipitations, recombinant protein expression and purification, *in vitro* SUMO assays, western blotting, structural analyses, immunofluorescent microscopy and analysis); M.J. (Protein expression and purification, NT647-labelling of WT E1 or K164Q-SAE2 E1, MST data analysis); A.J.G (siRNA and primer design, site-directed mutagenesis); P.W. and B.F.C. (MST sample measurement using Monolith NT.115 [NT.115Pico/N.T.LabelFree] instrument (NanoTemperTechnologies). T.J.K supervised P.W and B.F.C. J.R.M. supervised the project and wrote the paper. All authors wrote and edited the manuscript.

## Supplemental Figures

**Supplemental Figure 1.**
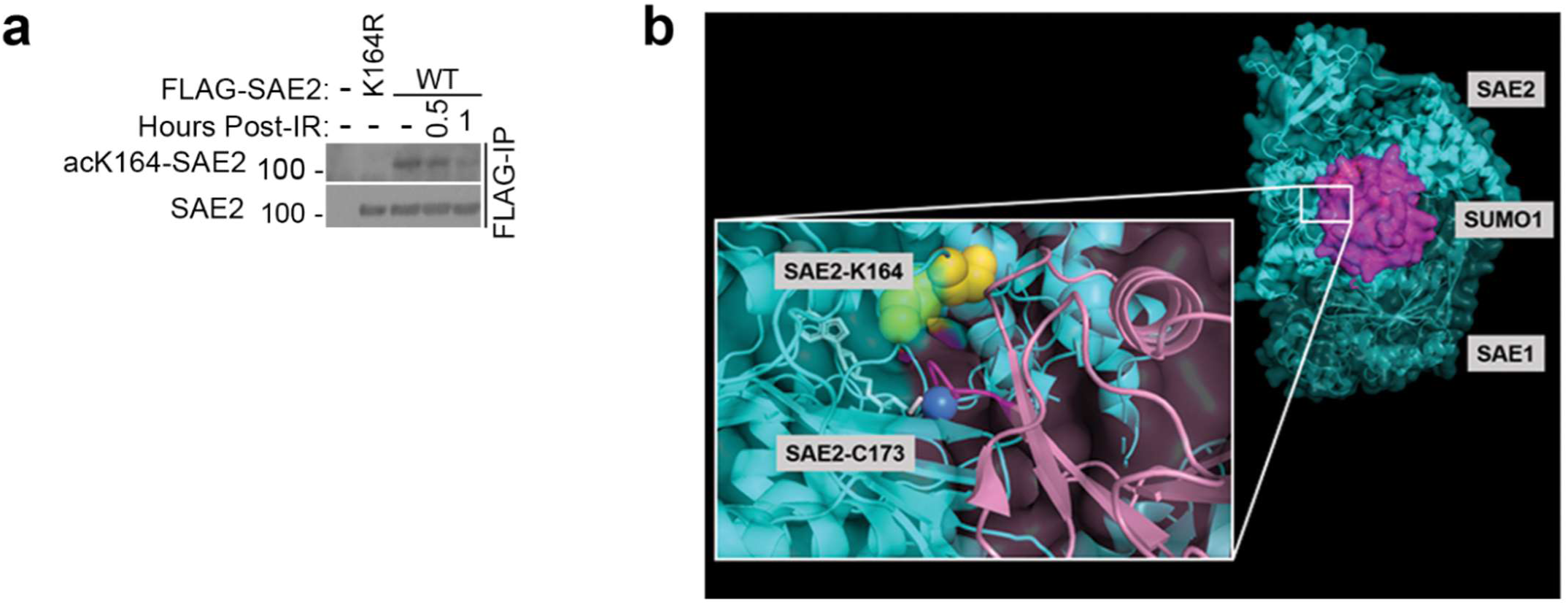
**a.** Western blot analysis of U2OS cells, untransfected (-) or expressing FLAG-SAE2 or FLAG-SAE2-K164R and treated with 10 Gy IR, then with half an hour (0.5) or 1 hour recovery (1). Lysates were subjected to anti-FLAG immunoprecipitation and western blots were probed with acetyl-K164-SAE2 (mouse monoclonal) and anti-SAE2 antibodies. **b.** Structure of SAE1:SAE2:SUMO1 (PDB: 3KYD) adapted from Olsen *et al.* (2010) represented as a ribbon structure of SAE1:SAE2 in cyan and SUMO1 in magenta. The magnified image shows the C-terminal tail of SUMO1 (dark magenta) extending toward SAE2-C173 (dark blue sphere) through a channel in SAE1:SAE2; the ceiling of the channel is in part formed by SAE2-K164 (yellow spheres).

**Supplemental Figure 2.**
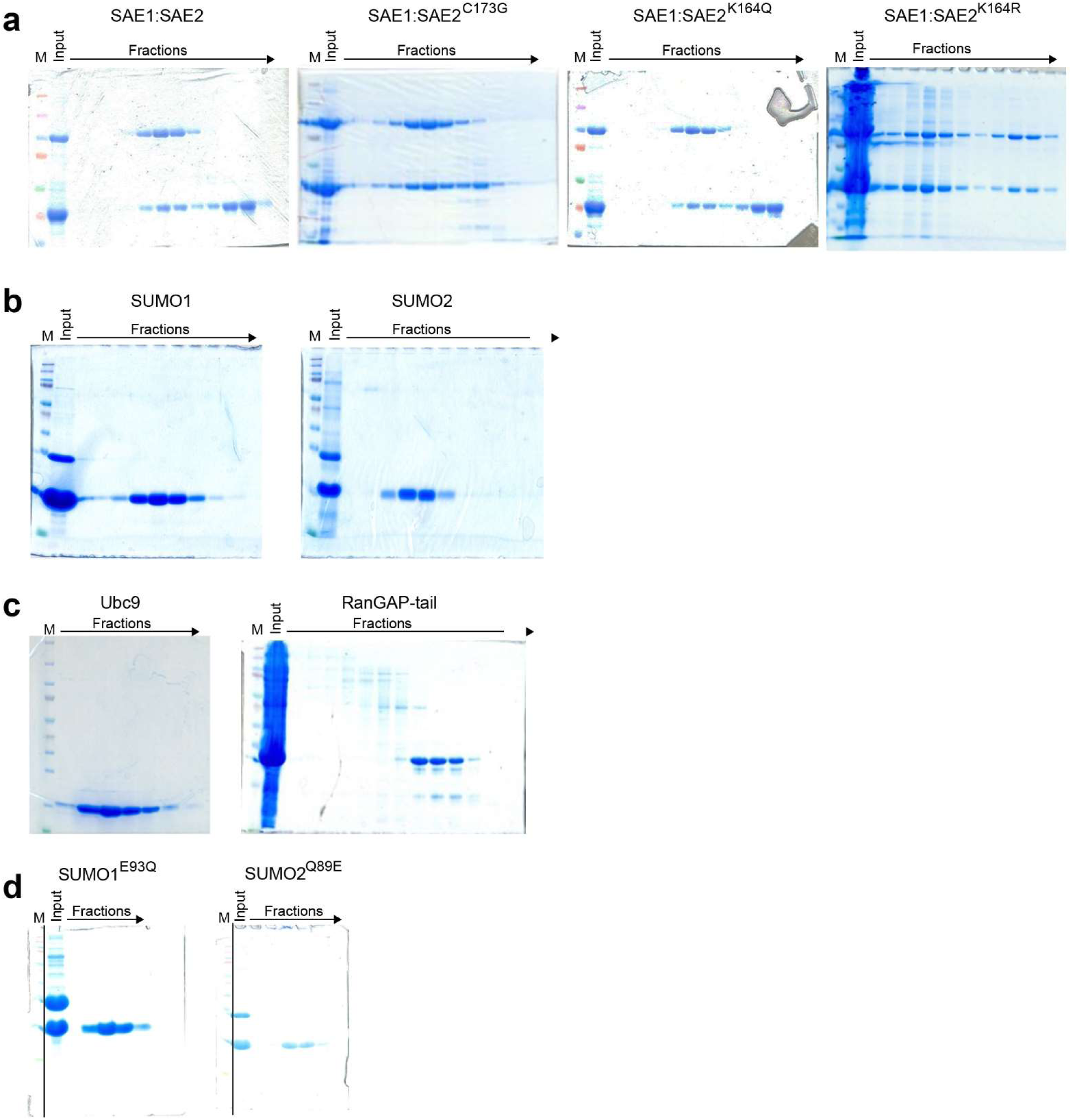
Coomassie gels for each recombinant protein prepared. Representative Instantblue stained SDS-PAGE gels from SEC fractions for the respective purified proteins. Shown here are gels from (**a**) SAE1:SAE2, SAE1:SAE2-C173G, SAE1:SAE2-K164Q, and SAE1:SAE2-K164R; (**b**) SUMO1 and SUMO2; (**c**) UBC9 and RanGAP1 (aa398-587); and (**d**) SUMO1-E93Q and SUMO2-Q89E protein purifications.

**Supplemental Figure 3.**
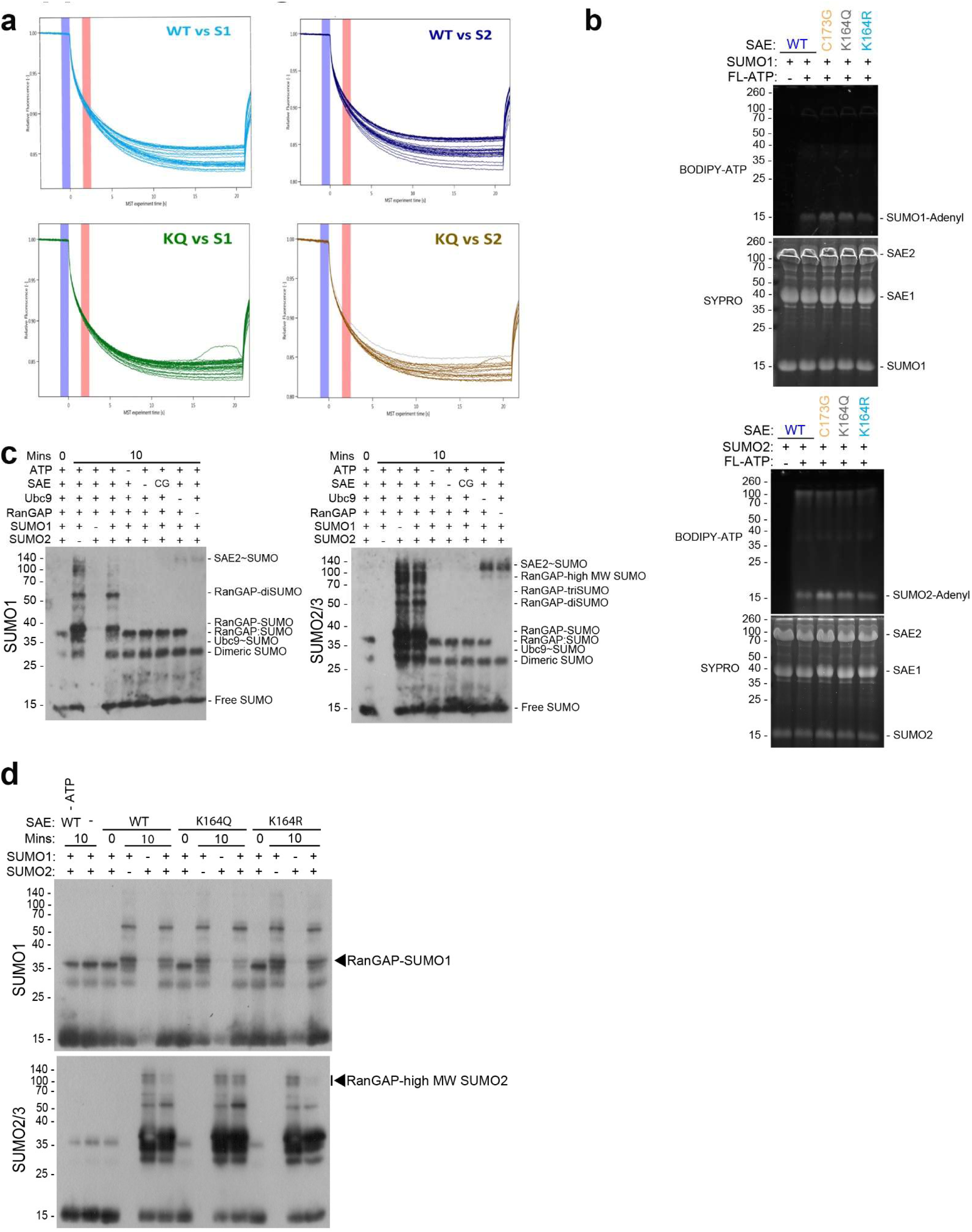
Extended *in vitro* data. **a.** MST Thermographs of SUMO1 or SUMO2 binding to SAE1:SAE2 or SAE1:SAE2-K164Q provide well-defined curves. The cold region is set to 0 s (blue) and the hot region set to 2.5 s (red) to determine the *K_d_* of the interaction and to avoid any potential convection phenomena. **b.** SDS-PAGE gels from *in vitro* adenylation assays after combining 30 μM SAE1:SAE2 variants, 40 μM SUMO1 (top) or SUMO2 (bottom), and 150 μM BODIPY-ATP. Gels were imaged with excitation at 488 nm to observe the BODIPY-ATP. Gels were subsequently stained using SYPRO ruby to check protein loading. **c.** *In vitro* RanGAP1 (aa398-587) SUMOylation control blots for SUMO1 and SUMO2/3. The first condition contains all reaction components but was not incubated at 30°C and gives 3 distinct bands, taken as unconjugated SUMO (15 kDa), dimeric SUMO (30 kDa), RanGAP1 (aa398-587): SUMO (37 kDa). These bands persist on incubation at 30°C for 10 minutes in conditions which omit ATP, SAE1:SAE2, UBC9, and in the use of the catalytically inactive SAE1:SAE2-C173G. The 37 kDa band is absent in the omission of RanGAP1 in the αSUMO1 and αSUMO2/3 blots, indicative of a SUMO- E1-E2-independent RanGAP1:SUMO species. SAE2∼SUMO is dependent on ATP, SAE1:SAE2 with catalytic activity and, therefore, accumulates in conditions –UBC9 and –RanGAP1 (aa398-587). RanGAP1-SUMO formation is dependent on all SUMOylation assay components, including incubation at 30°C, as is the formation of RanGAP1-polySUMO, which is most prominent in the αSUMO2/3 blot. Omission of SUMO protein variant controls abrogates the signal, indicating that the antibodies are SUMO variant specific. **d.** Representative blots for the *in vitro* SUMOylation data in Fig. 2d. Reactions comprised 20 μM SUMO1 and/or 20 μM SUMO2, 200 nM SAE1:SAE2, 500 nM UBC9, 10 μM RanGAP1 (aa398-587), and 5 mM ATP, incubated at 30°C for 10 minutes. Conditions were processed by SDS-PAGE in duplicate such that western blots were developed using αSUMO1 and αSUMO2/3 antibodies. Note that the bands indicated are those quantified.

**Supplemental Figure 4.**
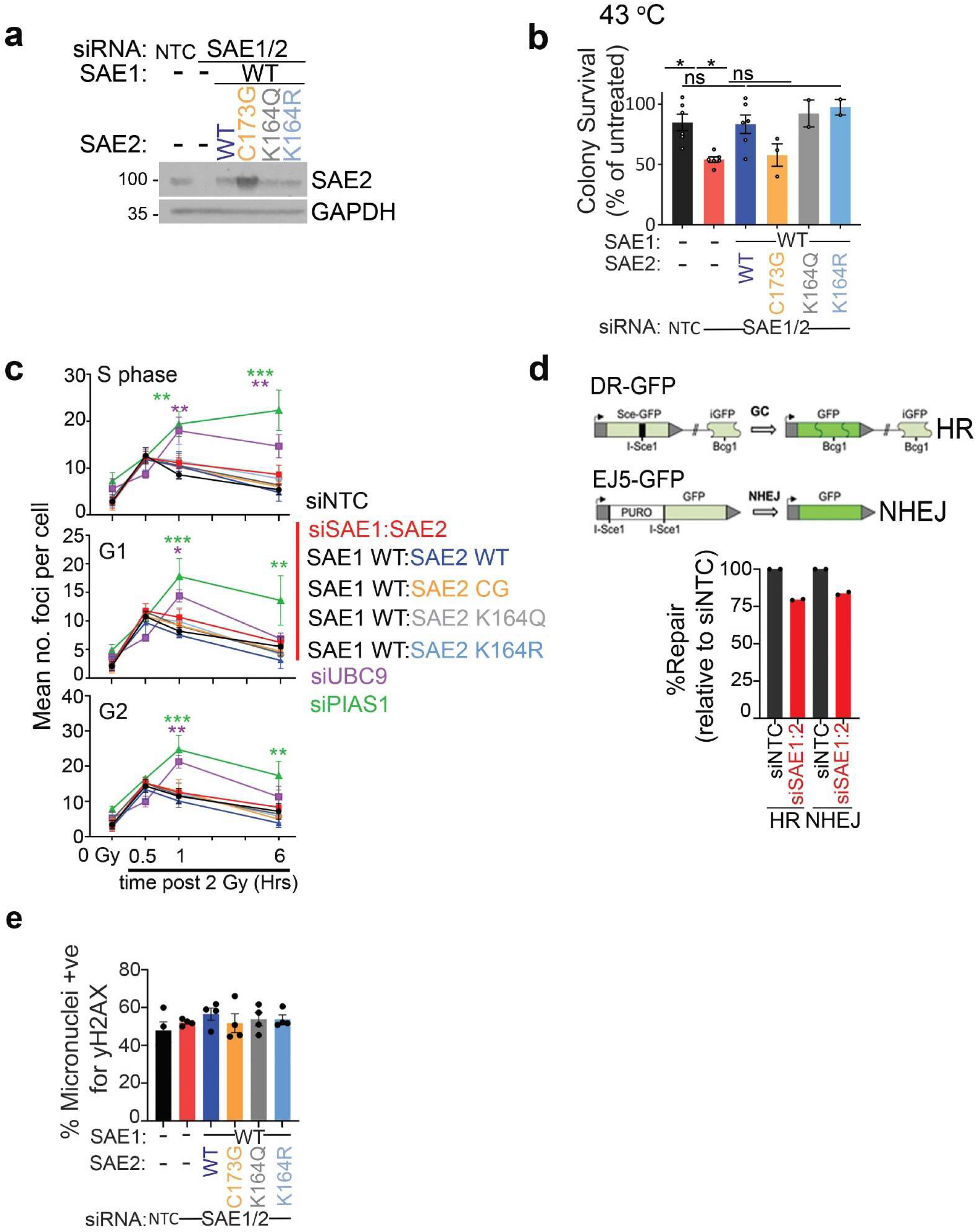
**a.** Representative western blot of SAE2 expression in stable inducible siRNA-resistant SAE1:SAE2 variant cells. Cells were treated for 72 hours with siRNA for either NTC or SAE1:SAE2 with the concurrent addition of 4 μg/ml Doxycycline to induce expression of the indicated integrated SAE1:2 constructs. **b.** U2OS depleted for SAE1:SAE2 and complemented with WT or indicated SAE1:SAE2-variant, subjected to 43°C for 40 minutes before replating and counting after colony growth. Significance calculated using one-way ANOVA. Error bars = SEM; N = 3. * = p≤ 0.05, ns = not significant p> 0.05. **c.** Automated analysis of γH2AX foci numbers, obtained through high-content microscopy, in cells treated with indicated siRNAs (siNTC- black, siSAE1:SAE2- red) with or without the complementation of inducible siRNA-resistant SAE2 variants (SAE2 WT- dark blue, SAE2 CG- orange, SAE2 K164Q- grey, SAE2 K164- light blue). siUBC9 (purple) and siPIAS1 (green) are used for comparison. Results displayed for data isolated from S phase (top), G1 (middle) and G2 (bottom) cell populations. Plotted data is derived from the mean number of foci per condition from 3 independent biological repeats. N = 3, error bars = SEM. Statistical significance was calculated using two-way ANOVA using Dunnett’s multiple comparisons test. Timepoints where there is a significant difference from the non-target control siRNA condition are marked with * = p<0.05, ** = p<0.01, *** = p<0.001. Purple and green * show that only siUBC9 and siPIAS1 conditions significantly deviate from siNTC at points in the time course. The total number of cells analysed for each data point is contained in Supp. Table 7. **d.** The measure of DNA repair from U2OS cells bearing integrated DNA repair reporter in cells treated with siNTC or siSAE1:siSAE2 and transfected with the enzyme, I-SCE-1. Illustration of the integrated DNA repair substrates for homologous recombination and non-homologous end-joining (Top). The graph (Bottom) displays the percentage of GFP-positive cells normalized to RFP-transfection efficiency. %-repair of siSAE1:SAE2 is given relative to siNTC. N = 2. Data from 2 independent biological repeats. f. The percentage of micronuclei positive for γH2AX in asynchronous siRNA-resistant SAE2 variant cells. Error bars SEM. Data from 4 independent biological repeats. n>50 micronuclei per condition per biological repeat. Significance tested using one-way ANOVA no significant differences between conditions identified.

**Supplemental Figure 5.**
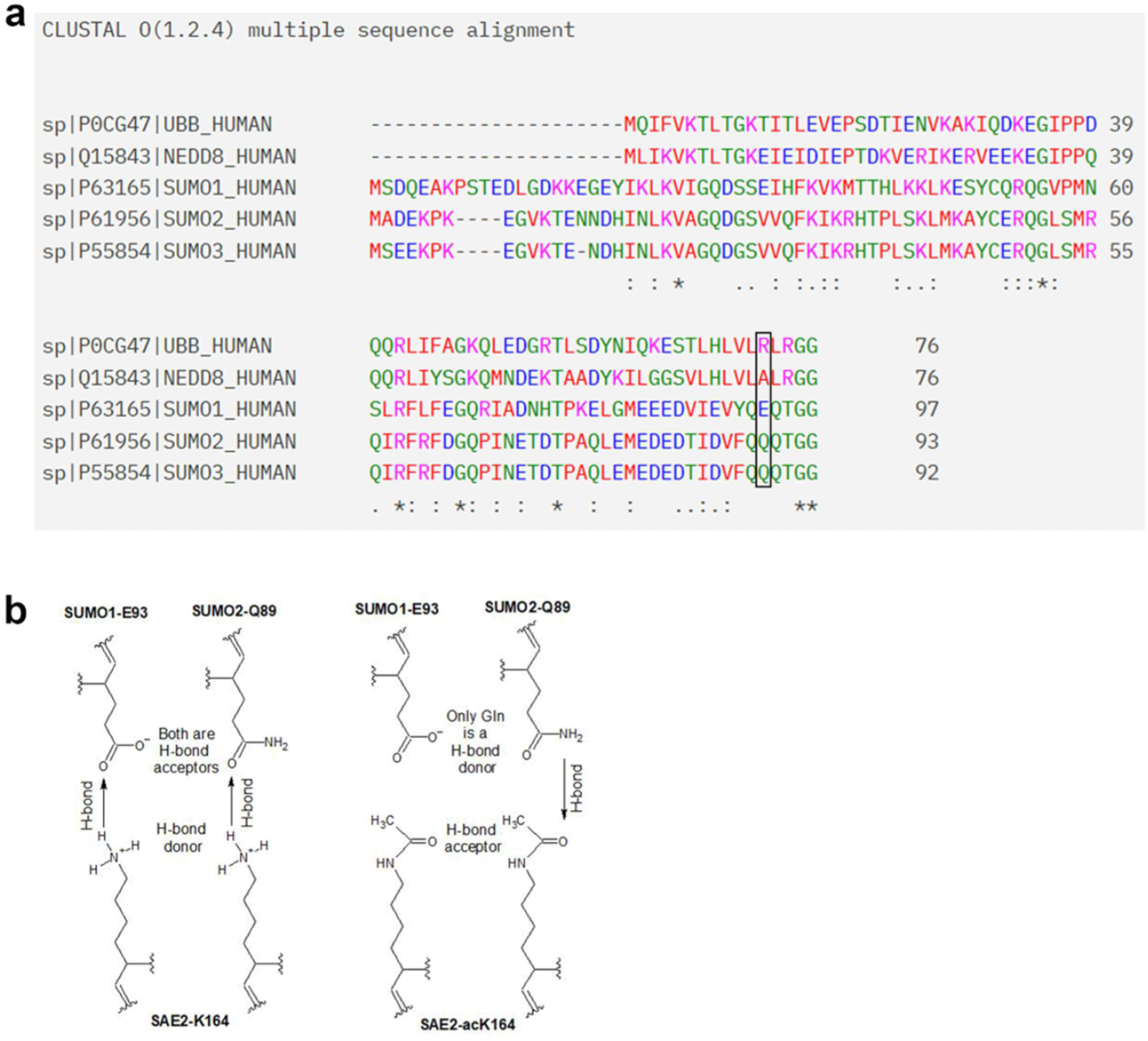
**a.** Amino acid sequence alignment for Ubiquitin (UBB), Nedd8, SUMO1, SUMO2, and SUMO3 up to the C-terminal di-Gly motif representative of mature activation/conjugation-competent Ubls. Black box indicates Ub-R72 which is divergent in Nedd8, SUMO1, and SUMO2/3 and required for Ubl E1 discrimination of different Ubl modifiers (Walden *et* al., 2003)[69]. Generated using Uniprot and Clustal Omega. **b.** Illustration of hypothetical proposed mechanisms of SAE1:SAE2-acK164 bias for SUMO2-Q89 over SUMO1-E93 through precise hydrogen bonding patterns. Unacetylated SAE2-K164 acts as a hydrogen bond donor to both SUMO1- E93 or SUMO2-Q89, SAE2-acK164 is exclusively a hydrogen bond acceptor which SUMO2-Q89 can act as a hydrogen bond donor to, while SUMO1-E93 cannot. This hypothetical hydrogen bonding arrangement explains why SAE1:SAE2- acK164 bears a bias towards SUMO2 activation.

## Materials and Methods

### acK164-SAE2 Antibody Generation

Custom mouse monoclonal (clone 30E2-2) was raised against acetylated K164- SAE2 peptide (HP[Lys-Ac]PTQRTFPGC) by GenScript. Available on request to the corresponding author subject to completion of an M.T.A.

### Generation of plasmids

See Supp. Table 1 for a comprehensive list of the vectors used. The SAE2-T2A-SAE2 pcDNA5/FRT/TO construct was designed by AJG and generated by GenScript using KpnI and NotI restriction sites. pET28a-SAE1 was cloned using NheI and BamHI restriction sites; pET28b-SAE2 was cloned using NcoI and NheI; pET23a-UBC9 was cloned using NdeI and BamHI; and pET23a-RanGAP1 was made using NdeI and BamHI restriction sites. pGEX4T-1-SUMO1-3 was designed by AJG and made by GenScript by cloning SUMO1-3 cDNA into BamHI and EcoR1 restriction sites. GST-NuMA constructs were generously gifted by Dr. Choi.

### Site-Directed Mutagenesis

Primers were designed for mutagenesis (Supp. Table 3), with mutagenesis performed by PCR using PfU (Promega). All mutagenesis was confirmed by Sanger Sequencing (Source Bioscience).

### Tissue culture

Parental FlpIn™ U2OS cells were obtained from Morris Lab cells stocks and grown in Dulbecco’s Modified Eagle Media (DMEM) supplemented with 10% Fetal Calf Serum (FCS) and 1% Penicillin/Streptomycin. Cells were cultured in Corning T75 flasks and 10 cm^2^ plates and kept at 37°C and 5% CO_2_. Once cells reached 70-80% confluency, they were passaged. Cells were tested for Mycoplasma by Hoescht staining. Cell lines have not been authenticated.

### Inducible stable cell line generation

U2OS^TrEx^-Flp-In™ (a gift from Grant Stewart, University of Birmingham) were co-transfected with SAE2-T2A-SAE1 cDNA in the pcDNA5/FRT/TO vector and the Flp-recombinase cDNA in the pOG44 vector at a 6:1 pcDNA5/FRT/TO DNA: pOGG44 DNA ratio using FuGene6 (Roche) at a ratio of 3.5:1 FuGene (μl): DNA (μg). Blank control transfections were performed as a control for selection. Two days after transfection, cells were selected with 150 μg/ml Hygromycin (Thermo Fisher Scientific) with culture medium replaced every 2-3 days; selection exerted for ∼2 weeks. After selection, cells were expanded and tested for expression of siRNA-resistant HA-SAE1/FLAG- SAE2 through treatment with siSAE1/siSAE2 (5 nM each) and 4 μg/ml Doxycycline for 72-hours. Cell lysates were prepared in 4xSDS loading buffer and western blot analysis performed.

### Plasmid and siRNA transfection

FuGene6 (Roche) was used at 2:1 FuGene (μl): DNA (μg), following the manufacturer’s guidelines. SUMO2 and SUMO1 overexpression was achieved using 0.5 µg of DNA per well of a 24 well plate for the durations indicated in the figure. GFP-NuMA constructs were transfected at 1 µg/ml. siRNA was introduced to cells using the transfection reagent Dharmafect1 (Dharmacon) following the manufacturer’s instructions. For a full list of siRNA sequences, see Supp. Table 4.

### FLAG immunoprecipitation

U2OS were cultured at 37°C/5% CO_2_ in 15 cm^2^ dishes supplemented with 4 μg/ml Doxycycline (Sigma) for 48 hours to induce exogenous SAE1:SAE2 expression. Cells washed with 1 ml ice-cold TBS (20 mM Tris/HCl pH 7.5, 150 mM NaCl) before suspension in RIPA Buffer (50 mM Tris-HCl pH 7.5, 150 mM NaCl, 1% TritonX100, 0.25% Sodium deoxycholate, 0.1% SDS, 1 mM EDTA, 10 mM NaF) plus EDTA-free protease inhibitor cocktail (Roche) and PhosSTOP (Roche). Lysis mix was incubated on ice for 10min and sonicated at 50% intensity for 10s. Samples were centrifuged at 14,000xg/4°C for 10 minutes, and supernatant combined with 15 μl FLAG(M2) agarose (Sigma), incubated with mixing O/N at 4°C. Beads were pelleted by centrifugation at 1,000xg/4°C for 2min. Supernatant was discarded and beads washed with TBST. 30 μl 4xSDS loading buffer was added to beads with boiling at 95°C for 10 minutes and centrifuged at 5,000xg/RT to pellet beads. Immunoprecipitations using histone acetyltransferase (HAT) inhibitors were conducted as above; however, before lysing, cells were incubated with 2.5 µM each of indicated inhibitor against p300 (A-485), TIP60 (NU9056), NAT10 (Remodelin hydrobromide) and GCN5 (Butyrolactone 3). To immunoprecipitate material from mitotic cells, 100 ng/ml of nocodazole was added for a total of 17 hours to relevant dishes, 48 hours after SAE1:SAE2 induction. HDAC inhibitors were applied at 2.5 µM for 2 hours prior to the release from Nocodazole treatment. At 17 hours, cells were washed twice with PBS and released into mitosis for 10 minutes in media supplemented with relevant HDAC inhibitor. Cells were then harvested via mitotic shake off and lysed as above in ice-cold RIPA Buffer (50 mM Tris-HCl pH 7.5, 150 mM NaCl, 1% TritonX100, 0.25% Sodium deoxycholate, 0.1% SDS, 1 mM EDTA, 10 mM NaF) plus EDTA-free protease inhibitor cocktail (Roche) and PhosSTOP (Roche) supplemented with 2 µM Panobinostat.

### Western blotting

For a list of antibodies, see Supp. Table 6. Protein samples in loading buffer were subject to SDS-PAGE and transferred onto Immobilon-P PVDF-membrane (Merck). Membranes were blocked in 5% milk in PBST or in 5 % BSA with TBST for 30 minutes. Incubation with primary antibodies for 16 hours/4°C/rolling. Membranes washed in PBST/TBST for 3×10 minutes and then incubated with relevant secondary HRP antibodies (Supp. Table 6) in blocking solution for minimum 1 hour/RT. Membranes washed in PBST/TBST for 3×10minutes and HRP stimulated with EZ-ECL mix (Biological Industries) or ECL Prime (Amersham). Blots exposed to X-ray film (Wolflabs) and developed with KONICA MINOLTA SRX-101A. Densitometry calculations performed using Image J.

### Protein overexpression and purification

BL21 (DE3; NEB) were transformed with the relevant plasmids (Supp. Table 1). Starter cultures were established by inoculating 40 ml LB (Melford; Kanamycin 50 µg/ml or Ampicillin 100 µg/ml) with a single colony, grown O/N at 37°C. 10 ml starter culture was used to inoculate each litre LB (Kan 50 µg/ml or Amp 100 µg/ml) and grown at 37°C/180 rpm to OD_595_ ∼0.6. Protein overexpression induced with (isopropyl-β-D-thiogalactoside) IPTG and temperature adjusted as follows, SUMO1 and SUMO2 at 0.5mM IPTG/18°C/18 hours; SAE1 and SAE2 at 1 mM IPTG/25°C/6 hours; UBC9 at 1 mM IPTG/37°C/4 hours; RanGAP1 (aa398-587) at 1 mM IPTG/37°C/4 hours; and FAT10 at 0.1 mM IPTG/25°C/5 hours, with shaking at 180 rpm. Overexpression protocols for SAE1:SAE2, UBC9, and RanGAP1 (aa398-587) were adapted from Flotho *et al.*, 2012 [63] and for FAT10 [79]. BL21(DE3) cells were harvested by centrifugation at 5,000xg/4 °C/10 minutes with the resulting pellet resuspended in 10 ml cold lysis buffer (20 mM Tris-HCl pH8, 130 mM NaCl, 1mM EGTA, 1mM EDTA, 1% Tritonx100, 10% Glycerol, 1 mM DTT, EDTA-free protease inhibitor (Roche)). Separately overexpressed SAE1 and SAE2 combined here. 0.5 mg/ml lysozyme was added and incubated for 30 minutes/4°C/rolling. 1 U/ml DNase (Thermo Fisher) added before sonication at 5×30 seconds at 100% intensity with 2-minute recovery – all on ice. Samples were centrifuged at 48,000xg/4°C/30 minutes in a JLA-25.50 and supernatant was filtered through a 0.45 μm PES membrane (Millex) and combined protein-tag-specific resin. Prior to use, resins were washed twice in PBS and once in the lysis buffer, with centrifugation performed at 1,000xg/4°C/3 minutes. Respective protein purification continued as follows.

### SUMO1 and SUMO2 purification

GST-SUMO1/SUMO2 were combined with 250 μl glutathione Sepharose 4B beads (Cytiva) and incubated for 3 hours/4 °C/rolling. Beads were centrifuged at 1,000xg/4°C/10 minutes with supernatant collected. Wash steps comprised 3×10 ml lysis buffer suspension of beads and 1×10 ml cleavage buffer (20 mM Tris-HCl pH 8.4, 150 mM NaCl, 1.5 mM CaCl_2_) with centrifugation as above. Beads were then suspended in 500 μl cleavage buffer supplemented with 16 U Thrombin cleaving protease and incubated at 4°C/16 hours/rolling. Samples were then centrifuged at 1,000xg/4°C/3 minutes and the supernatant was collected for centrifugation at 14,000xg/4°C/20 minutes to clear any beads or aggregate. The 500 μl sample was subjected to size-exclusion chromatography (SEC) through an AKTA pure™ (UNICORN™ software) Superdex200 Increase 10/300 GL column equilibrated in 20 mM Hepes pH 7.5, 100 mM NaCl, 0.5 mM TCEP: 0.5 ml fractions collected. Fractions constituting a UV_280_ trace peak were analysed by SDS-PAGE stained with InstantBlue (Lubioscience; Supp. Fig. 2b-e). Pure protein fractions were pooled and stored at -80°C. See Supp. Table 2.

### UBC9 purification

Column filled with 10 ml SP-Sepharose beads and 60 ml UBC9 lysate applied at ∼1 ml/min; FT collected and reapplied. UBC9 lysis buffer was passed through a column for wash step. 20 ml UBC9 elution buffer (50 mM Na-phosphate pH 6.5, 300 mM NaCl, 1 mM DTT, 1 cOmplete protease inhibitor (EDTA-free) tablet/50 ml) applied to beads and 1.5 ml fractions collected - 10 μl samples analysed by 15% SDS-PAGE and InstantBlue stain. Fractions with the greatest quantity and purity of UBC9 protein combined and concentrated down to 5 ml using 3-kDa MWCO centrifugal concentrator (Thermo Scientific) at 4,000xg/4 °C. Sample cleared by centrifugation at 14,000xg/4°C/20 minutes before SEC through a Superdex75 equilibrated in transport buffer (20 mM Hepes pH 7.3, 110 mM potassium acetate, 1 mM EGTA, 1 mM DTT, 1 cOmplete protease inhibitor tablet/L): 4 ml fractions collected. Fractions constituting UV_240_ peak were analysed by 15% SDS-PAGE and Instantblue stain. Pure UBC9 protein fractions (Supp. Fig. 2e) were pooled and concentrated as before, and finally aliquoted and stored at -80°C. See Supp. Table 2

### SAE1:SAE2 and RanGAP1 (aa398-587) purification

These His-tagged protein lysates were combined with 1 ml nickel beads (Sigma) and incubated at 4°C/2 hours/rolling. Samples were then centrifuged at 1,000xg/4°C/10 minutes to pellet nickel beads; FT collected and retained. Nickel-bead:His-protein pellet suspended in 10 ml wash buffer ahead of centrifugation as before; supernatant retained. Nickel beads resuspended in 5 ml elution buffer and centrifuged as before; supernatant extracted and pushed through 0.45 μm PES filter. This protein suspension was run using an AKTA pure™ (UNICORN™ software) on a HiLoad 16/600 Superdex200 pg column equilibrated with 20 mM Hepes pH 7.5, 100 mM NaCl, 0.5 mM TCEP buffer: 2 ml fractions collected. Fractions corresponding with a UV_280_ peak were analysed by SDS-PAGE stained with InstantBlue. Fractions containing the purest SAE1:SAE2 or RanGAP1 (aa398-587; Supp. Fig. 2a & e) were pooled and exposed to 4,000xg at 4°C in centrifugal concentrators with 30 kDa and 10 kDa MWCO, respectively. Proteins were aliquoted and stored at -80°C. See Supp. Table 2.

### Microscale thermophoresis (MST)

SAE1:SAE2 was labelled using Protein Labelling Kit RED-NHS 2nd Generation (NanoTemper Technologies). The labelling reaction was performed according to the manufacturer’s instructions in the supplied labelling buffer, using 20 μM SAE1:SAE2 and a molar dye: protein ratio ≈ 3:1 at RT for 30 minutes in the dark. Unreacted dye was removed with the supplied dye removal column equilibrated with MST buffer (20 mM Hepes pH 8.35, 150 mM NaCl, 0.5 mM TCEP). The degree of labelling was determined using UV/VIS spectrophotometry at 650 and 280 nm. A degree of labelling of 0.5-0.6 was typically achieved.

The labelled SAE1:SAE2 protein was adjusted to 20 nM with MST buffer supplemented with 0.005% Tween20. The SUMO1/SUMO2 ligand was dissolved in MST buffer supplemented with 0.005% Tween20, and a series of 16 1:1 dilutions was prepared using the same buffer. For the measurement, each ligand dilution was mixed with one volume of labelled SAE1:SAE2 protein for a final concentration of 10 nM and ligand concentrations ranging from 125 mM to 0.00381 μM, respectively. After 10 minutes incubation, samples were centrifuged at 10,000xg for 10 minutes, and loaded into Monolith NT.115 [Premium] Capillaries (NanoTemper Technologies). MST was measured using a Monolith NT.115 [NT.115Pico/N.T.LabelFree] instrument (NanoTemper Technologies) at an ambient temperature of 22°C. Instrument parameters were adjusted to 20% LED power and high [low/medium] MST power. Data of three independently pipetted measurements were analysed (MO.Affinity Analysis software version 2.3, NanoTemper Technologies) using the signal from an MST-on time of 2.5s.

### SUMO adenylation assay

Reactions were performed by combining 30 µM SAE1:SAE2-K164 variants with 40 µM SUMO1, and 1 U pyrophosphatase. Reactions were initiated by adding 150 µM BODIPY-ATP and incubated at 30°C for 10 minutes. Reactions were quenched by the addition of 4xloading buffer and incubated at 95°C for 5 minutes before running samples on 15% SDS-PAGE. Gels were imaged using excitation at 488 nm to excite BODIPY-ATP and bands at 15 kDa were taken to be SUMO1-AMP-BODIPY.

### SAE1:SAE2-loading assays

Conducted in V_T_ 20 μl in SAB with 5 μM SAE1:SAE2 and 5 mM ATP with reactions started by adding 10 μM SUMO1-C52A-S9C-Alexa488 or SUMO2-C48A-A2C-Alexa64 and incubating samples on ice for 15 seconds. The reaction was terminated with 20 μl 4xLoading buffer (reducing-agent free) and incubated at 95°C. Samples were processed for analysis by SDS-PAGE and imaged at excitation wavelengths of 488 nm and 647 nm, respectively. The at 120 kDa was taken as SAE2∼SUMO product.

### SUMOylation assays

All proteins were diluted in SUMO assay buffer (SAB; 20 mM Hepes pH7.5, 50 mM NaCl, 5 mM MgCl_2_) and quantified on a NanoDrop2000/2000c (Thermo Fisher Scientific), using SAB to adjust proteins to target concentrations as in Supp. Table 2. Reaction mixes prepared to V_T_ 20 μl SAB with 200 nM SAE1:SAE2, 500 nM UBC9, 10 μM RanGAP1 (aa398-587), 20 μM SUMO1, and 20 μM SUMO2. The reaction was started by adding 5 mM ATP with incubation at 30°C for 10 minutes, and reactions were terminated by adding 20 μl 4xLoading buffer (reducing-agent free) and incubating samples at 95°C for 10 minutes. Samples were centrifuged at 14,000xg for 5 minutes before processing by SDS-PAGE and western blot analysis.

### Densitometry

Densitometry was calculated using ImageJ [80] to quantify western blot band intensities. All quantification is from at least 3-independent experiments. To quantify RanGAP1-SUMO from *in vitro* assay western blots, band intensities were measured, and the background was subtracted. Values were normalised against the WT SAE1:SAE2:SUMO1/SUMO2-only condition RanGAP1-SUMO product intensity. For the densitometry to calculate the levels of SUMO conjugation, we calculated the relative amounts of SUMO ‘smear’ in cells using densitometry with Image J. The amount was then normalised to a GAPDH loading control.

### Immunofluorescent staining

The staining for γH2AX foci kinetics was performed as follows. Cells were plated directly onto 48 well plates at 1 x 10^4^ cells/ml and allowed to settle overnight. siRNA was applied to cells for 72 hours and complemented with SAE1:SAE2 variants by the addition of 4 μg/ml Doxycycline. Prior to 2 Gy irradiation cells were pulsed with 1 µM EdU for 10 minutes. 0 Gy samples were fixed directly after Edu incubation. After irradiation, cells were allowed to recover for allotted timepoints. At allotted timepoints cells were pre-extracted using CSK buffer (100 mM NaCl, 300 mM sucrose, 3 mM MgCl_2,_ 10 mM PIPES pH 6.8, 0.7% Triton x100) for 1 minute RT before fixation with 4 % PFA in PBS. Fixed cells were permeabilised for a further 5 minutes using 0.5% TritonX100 in PBS before incubation with blocking solution (10% FCS in PBST) for 30 minutes. EdU was labelled by Click-iT® chemistry according to the manufacturer’s protocols (Life Technologies) with Alexa-647-azide. Cells were washed then blocked for a further 30 minutes before incubation with primary antibody diluted in blocking solution for 1 hour RT. See Supp. Table 6 for a list of antibodies. Following this the samples were washed in PBST before incubation with the fluorescent secondary antibody for 1 hour RT. Samples were washed three times in PBS before the DNA was stained using Hoescht at a 1:50,000 concentration for 5 minutes. Excess of Hoescht was washed with PBS before antibodies were fixed in place for 5 min using 4% PFA. PBS was reapplied to cells and imaging proceeded within 3 days.

Cells for micronuclei and mitotic spindle staining were plated at 2 x 10^4^ cells/ml in 24 well plates on glass coverslips and treated with siRNA and doxycycline as described above. Micronuclei samples were fixed 72 hours later with 4% PFA before immunofluorescent staining.

Mitotic spindle samples were all treated with 100 ng/ml nocodazole for 16h. Cells were then washed twice in PBS before media was replaced to allow mitosis to progress for 35 minutes before fixation with PFA. Samples using siRNA resistant constructs were treated with siRNA and Doxycycline for 48h prior to the addition of nocodazole and cells were released into mitosis in media supplemented with Doxycycline. Deviations and additions to this basic protocol such as the use of inhibitors and or additional transfections are outlined schematically in the relevant figure.

Immunostaining of cells for micronuclei and mitotic spindle apparatus proceeded as follows: Cells were permeabilised for 5 minutes using 0.5% TritonX100 in PBS and blocked in blocking solution for 1 hour RT before the addition of primary antibodies. See Supp. Table 6 for a list of antibodies. Samples were washed in PBST before the addition of secondary antibodies. DNA was stained with Hoechst and coverslips were mounted onto glass slides.

### Microscopy and analysis

High content γH2AX foci imaging and analysis was conducted using the CellInsight CX5 HCS Platform with 20x objective lens using Compartmental Analysis BioApplication software. Spot detection within the nuclear compartment was used to count foci. Raw data outputs from this analysis were extracted so that cell cycle positioning could be achieved by plotting the total nuclear intensity of the Hoechst signal against the log average nuclear intensity of the EdU signal. This was used to separate γH2AX foci numbers into S phase, G1 and G2 populations. All other staining was imaged using the Leica DM6000B microscope using a 40x objective and HBO lamp with 100W mercury short arc UV bulb light source and four filter cubes, A4, L5, N3 and Y5, which produce excitations at wavelengths 360, 488, 555, and 647 nm, respectively. To analyse micronuclei, all cells were counted in at least 4 fields of view to reduce sampling bias, and a minimum of 400 cells were counted per condition per experimental repeat. Mitotic spindles were assessed by counting all laterally presented metaphase and anaphase cells in a field of view, with a minimum of 50 cells counted per condition per experimental repeat. All cells in a field of view were counted from least 4 fields of view. Metaphase and anaphase cells were identified based on the well characterised morphology that tubulin and DNA adopt during these mitotic stages, pericentrin staining was used as a further aid this determination when a spare channel for imaging allowed for its inclusion.

### Denaturing SUMO immunoprecipitation on mitotic cells

The methodology for denaturing SUMO immunoprecipitations was adapted from [12]. In brief, 8x 10cm dishes were plated per condition. At 80% confluency cells were treated with 100 ng/ml nocodazole, 1 µM ML792 and 4 μg/ml Doxycycline for 16 hours. Cells were washed twice in PBS, released into growth media supplemented with ML792 and Doxycycline for 10 minutes before harvesting via mitotic shake off. The resulting pellets were lysed in 200 µl 1% SDS lysis buffer (20 mM sodium phosphate, pH 7.4, 150 mM NaCl, 1% SDS, 1% Triton, 0.5% sodium deoxycholate, 5 mM EDTA, 5 mM EGTA, 10 mM NEM, plus EDTA-free protease inhibitor cocktail (Roche) and PhosSTOP (Roche) and sonicated until the viscous sample became fluid. Samples were boiled for 10min with 50mM DTT and centrifuged at 14,000g for 15 minutes. 30 µl of supernatant was taken and combined with 30 µl of 4xSDS loading buffer for use as an input on western blots. The remaining supernatant was diluted 1:10 in RIPA without SDS (20 mM sodium phosphate, pH 7.4, 150 mM NaCl, 1% Triton, 0.5% sodium deoxycholate, 5 mM EDTA, 5 mM EGTA, 10 mM NEM, plus EDTA-free protease inhibitor cocktail (Roche) and PhosSTOP (Roche).Total protein was quantified for each sample using Pierce™ 660 nm Protein Assay Reagent following manufacturer guidelines. The lysates were equalised for protein content and split so that half was used for SUMO1, and half was used for SUMO2/3 immunoprecipitations. Pierce™ Protein A/G Agarose beads (Thermo Fisher) were washed 3x in 0.1% SDS RIPA (20 mM sodium phosphate, pH 7.4, 150 mM NaCl, 0.1% SDS, 1% Triton, 0.5% sodium deoxycholate, 5 mM EDTA, 5 mM EGTA) before rotating in 0.1% RIPA with primary antibody at room temperature for 1 hour. Thirty microliters of beads were prepared for each sample using 3 µg of antibody per 30 µl of beads, ab32058 (Abcam) was used for SUMO1 and ab81371 (Abcam) was used for SUMO2/3 immunoprecipitations. Antibody-bound beads were pelleted and combined with prepared lysates prior to rotation at 4°C O/N. Beads were pelleted and washed 3x 3 mins with agitation in 0.1% RIPA before protein elution at 95°C in 4xSDS loading buffer.

### Colony survival assays

U2OS Flp-In™ cells were plated in 24-well plates at 2×10^4^/ml and treated with siSAE1:siSAE2 (5 nM each) and Dox for 72 hours. Cells were heat shocked in a water bath at 43°C for 40 minutes with control condition placed in a 37°C water bath. Cells were suspended in 100 μl 1xTrypsin followed by 900 μl DMEM and plated at limiting dilutions in 6-well plates. Plates were incubated for 7 days at 37°C and 5% CO_2,_ then stained with 0.5% crystal violet (50% methanol) and counting.

### DR-GFP and NHEJ-EJ5

U2OS-DR3-GFP and NHEJ-EJ5 (reporter cell lines) were a generous gift from Jeremy Stark (City of Hope, Duarte U.S.A.). U2OS reporter cell lines were simultaneously co-transfected with siRNA using Dharmafect1 (Dharmacon) and DNA (RFP and *I-Sce1* endonuclease expression constructs) using FuGene6 (Promega), respectively. After 16 hours, the media was replaced, and cells were grown for a further 48-hour before fixation in 2% PFA in PBS. RFP and GFP double-positive cells were scored by FACS analysis using a CyAn flow cytometer. Ten thousand cells were counted per sample. Data was analysed using Summit 6.2 software. The percentage of GFP-positive cells was determined as a fraction of RFP positive cells (RFP only +double GFP-RFP) to control for transfection efficiency.

### Statistics and Reproducibility

All statistics were done using two-sided unpaired t-test, one-way ANOVA or two-way ANOVA. Significance shown as *p<0.05, **p<0.01, ***p<.0.001, n.s = not-significant p>0.05. All experiments were repeated to generate biological replicates. The n-value is reported for each experiment.

## Supplemental Tables

**Supplemental Table 1.**
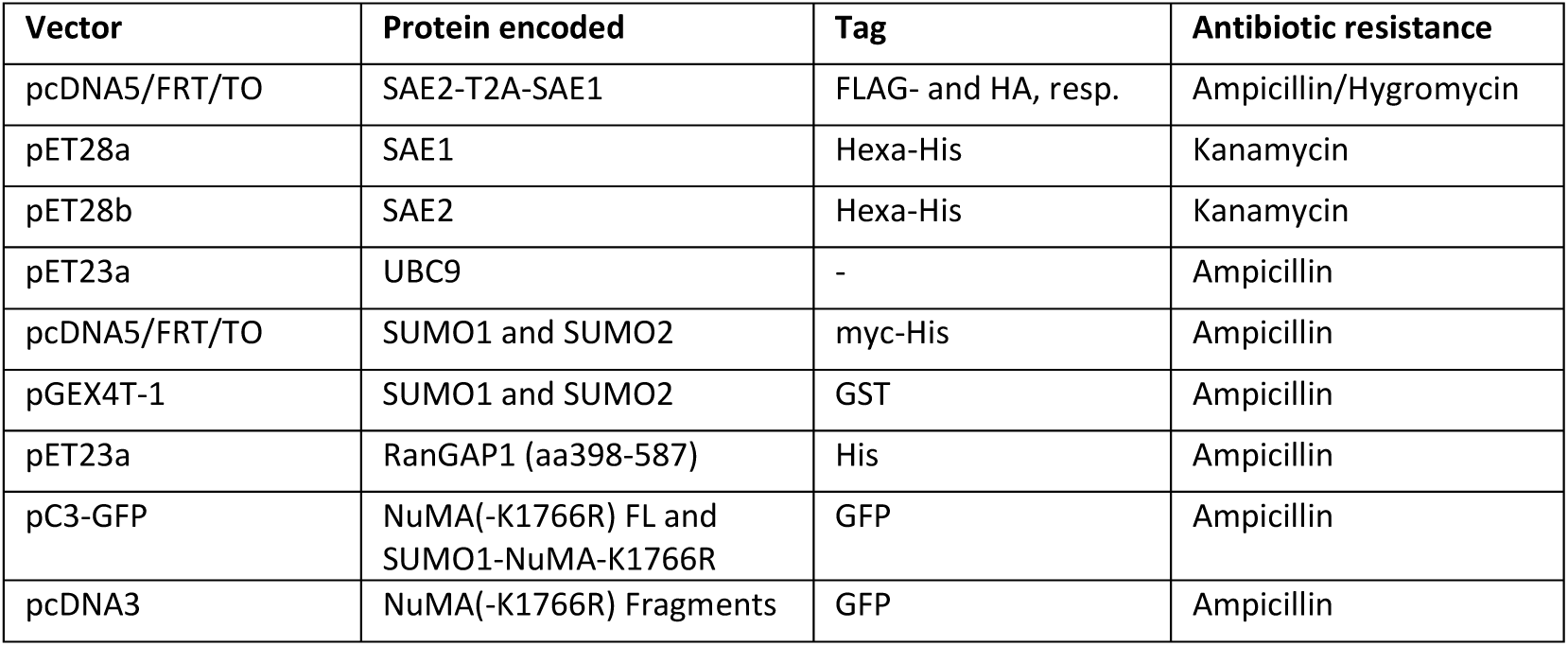
Vectors, gene(s) of interest encoded, and antibiotic resistance.

**Supplementary Table 2.** Recombinant proteins and their parameters.

| Protein | Molecular weight (kDa) | Extinction coefficient ($\epsilon/1000$)<br>-Cys reduced- | Useful target<br>concentration (mg/ml) |
| --- | --- | --- | --- |
| SAE1:SAE2 | 109.655 | 69.330 | 0.200 |
| UBC9 | 18.007 | 29.700 | 0.150 |
| RanGAP1(aa398-587) | 22.386 | 10.805 | 4.500 |
| SUMO1 | 11.277 | 4.470 | 2.000 |
| SUMO2 | 10.753 | 1.490 | 2.000 |

Useful protein preparation concentrations for appropriated pipetting volumes for *in vitro* reactions.

**Supplementary Table 3.**
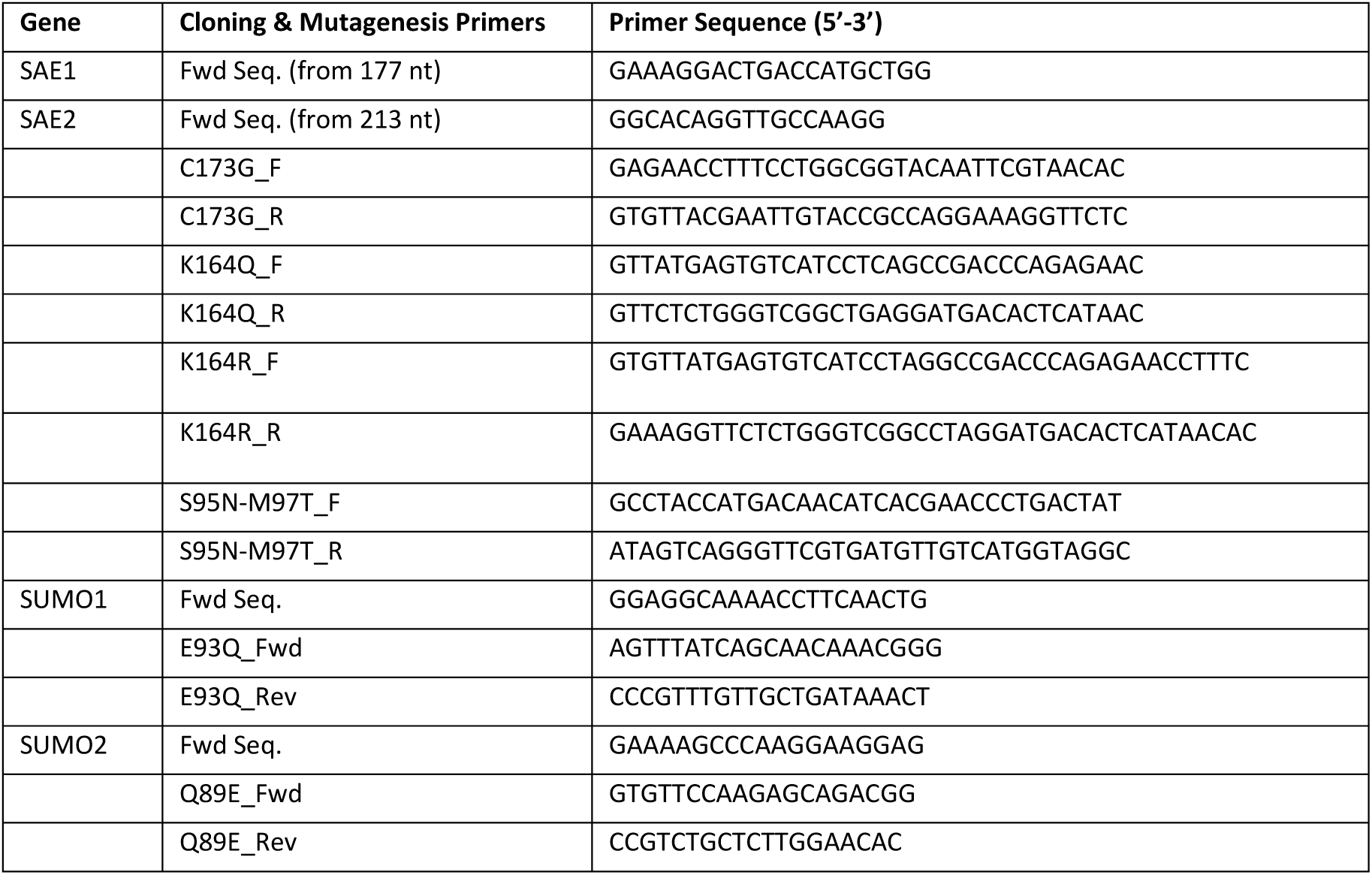
Oligonucleotide primers used for cloning and site-directed mutagenesis.

**Supplementary Table 4.**
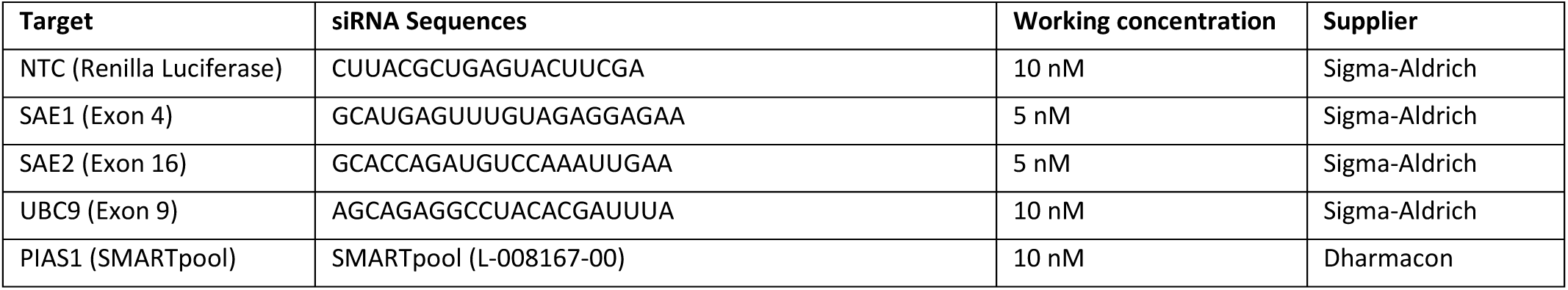
siRNA sequences.

**Supplementary Table 5.**
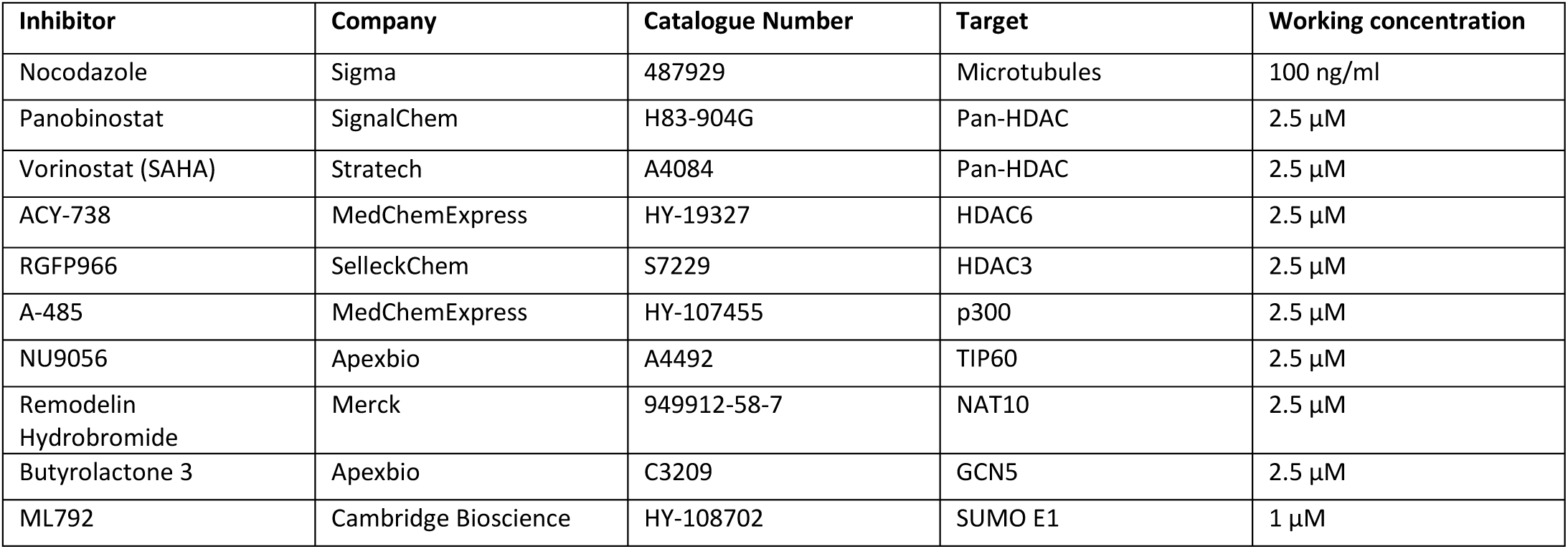
Details of inhibitors.

**Supplementary Table 6.** Antibodies.

| Antibody (clone) | Host | Supplier | Cat. number | Lot number | Use | Conc. | RRID |
| --- | --- | --- | --- | --- | --- | --- | --- |
| β-actin | Rabbit | Abcam | Ab8227 | GR3215935-1 | WB | 1:3000 | AB_2305186 |
| acK164-SAE2 (30E2-2) | Mouse | GenScript | Custom design | NA-6 | WB | 1:1000 | N/A |
| SAE1 | Rabbit | Abcam | ab185552 | GR3379016-1 | WB | 1:2500 | - |
| SAE2 | Rabbit | Sigma | HPA041436 | R38238 | WB | 1:2500 | AB_2677479 |
| UBC9 | Rabbit | Abcam | ab75854 | GR118836-6 | WB | 1:2500 | AB_1310787 |
| SUMO1 (Y299) | Rabbit | Abcam | ab32058 | GR3244068-3;<br>GR3366977-1;<br>GR3244068-3;<br>GR3366977-9; | WB/IP | 1:5000 | AB_778173 |
| SUMO1 (EP298) | Rabbit | Abcam | ab133352 | GR268526-16;<br>GR268526-12;<br>GR268526-10 | WB | 1:500 | AB_11156108 |
| SUMO2/3 (8A2) | Mouse | Abcam | ab81371 | GR3379145-7 | WB/IP | 1:1000 | AB_1658424 |
| His | Mouse | Sigma | H1029 | 025M4780V | WB | 1:5000 | AB_260015 |
| FLAG (M2) | Mouse | Sigma Aldrich | F1804 | SLBT7654 | WB | 1:1000 | AB_262044 |
| GFP | Mouse | Roche | 11814460001 | 47859600 | WB | 1:2500 | AB_390913 |
| γH2AX | Rabbit | Abcam | Ab2893 | GR3242597-1 | IF | 1:2000 | AB_303388 |
| αTubulin | Mouse | Santa Cruz | sc-5286 | H0613 | WB | 1:1000 | AB_628411 |
| CENPA | Mouse | Invitrogen | MA1-20832 | YL4140131 | IF | 1:500 | AB_2078763 |
| NuMA | Mouse | Santa Cruz | sc-365532 | A3124<br>C1323 | WB<br>IF | 1:1000<br>1:2000 | AB_10846197 |
| Vinculin [EPR8185] | Rabbit | Abcam | ab129002 | GR221671-50 | WB | 1:2000 | AB_11144129 |
| Pericentrin | Rabbit | Abcam | ab4448 | GR245491-2 | IF | 1:1000 | AB_304461 |
| αTubulin | Mouse | Novus | NB100-690 | G-3 | IF | 1:500 | AB_521686 |
| pS10-H3 | Mouse | Invitrogen | MA5-15220 | XJ3742465 | WB | 1:5000 | AB_11008586 |
| pS10-H3 | Rabbit | Antibodies.com | A94899 | 32699 | WB | 1:5000 | - |
| Donkey $\alpha$ Mouse AlexaFluor 488 | Donkey | Life Technologies | A21202 | 1975519 | IF | 1:5000 | AB_141607 |
| Donkey $\alpha$ Rabbit AlexaFluor 488 | Donkey | Life Technologies | A21206 | 1874771 | IF | 1:5000 | AB_2535792 |
| Donkey $\alpha$ Mouse AlexaFluor 555 | Donkey | Life Technologies | A31570 | 1774719 | IF | 1:5000 | AB_2536180 |
| Donkey $\alpha$ Rabbit AlexaFluor 555 | Donkey | Life Technologies | A31572 | 1945911 | IF | 1:5000 | AB_162543 |
| Donkey $\alpha$ rat AlexaFluor 555 | Donkey | Life Technologies | A21434 | 1987272 | IF | 1:5000 | AB_2535855 |
| Rabbit $\alpha$ Mouse HRP | Rabbit | Dako | P0161 | 20062080 | WB | 1:10000 | AB_2687969 |
| Swine $\alpha$ Rabbit HRP | Swine | Dako | P0217 | 20047666 | WB | 1:10000 | AB_2728719 |
| Mouse TrueBlot® ULTRA: Anti-Mouse Ig HRP | Rat | Rockland | 18-8817-30 | 39891 | WB | 1:5000 | AB_2610849 |

**Supplementary Table 7.**
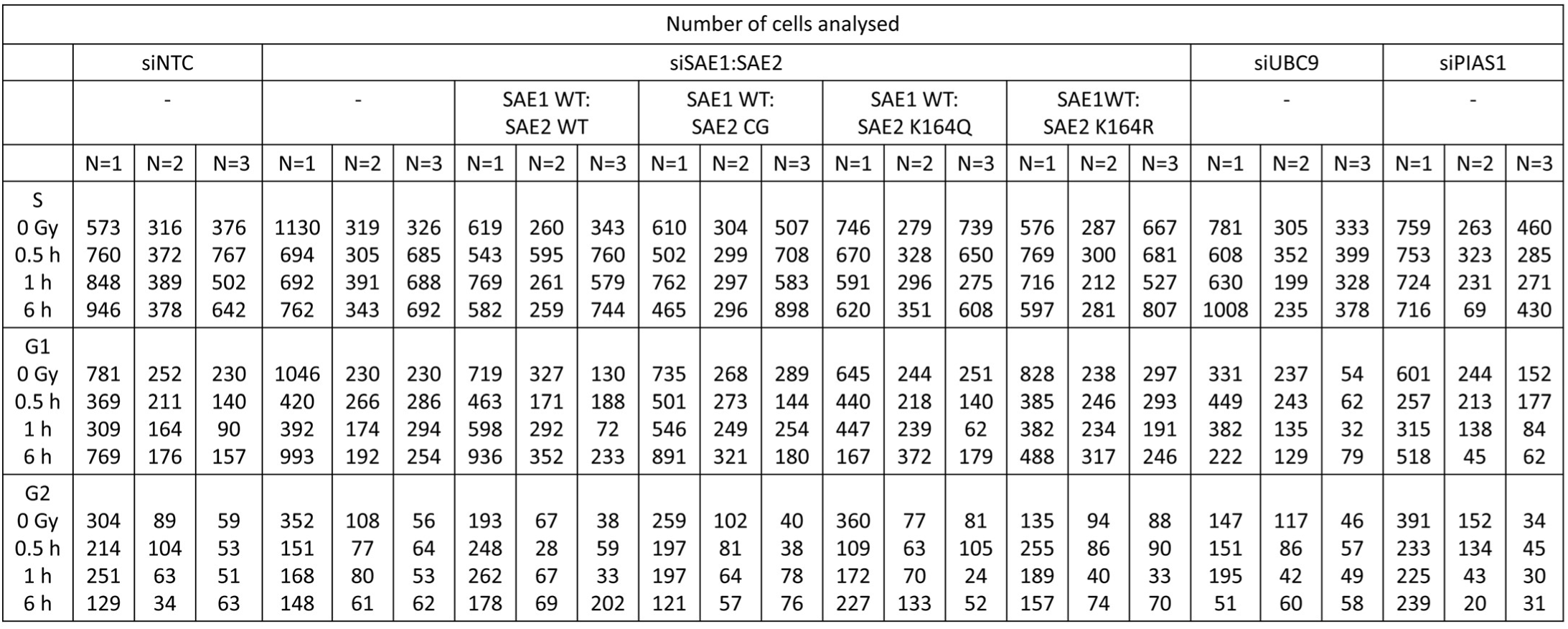
Number of cells analysed in automated analysis of γH2AX foci separated by cell cycle positioning.

## Notes

### Competing Interest Statement

The authors have declared no competing interest.

### Summary of Updates

In the update, we identify that a fusion of the mitotic SUMOylation substrate NuMA with SUMO1, can rescue spindle defects in cells treated with an HDAC6 inhibitor or complement with K164Q-SAE2.

